# Attentional Modulations of Alpha Power Are Sensitive to the Task-relevance of Auditory Spatial Information

**DOI:** 10.1101/2021.02.12.430942

**Authors:** Laura-Isabelle Klatt, Stephan Getzmann, Daniel Schneider

## Abstract

The topographical distribution of oscillatory power in the alpha band is known to vary depending on the current focus of spatial attention. Here, we investigated to what extend univariate and multivariate measures of post-stimulus alpha power are sensitive to the required spatial specificity of a task. To this end, we varied the perceptual load and the spatial demand in an auditory search paradigm. A centrally presented sound at the beginning of each trial indicated the to-be-localized target sound. This spatially unspecific pre-cue was followed by a sound array, containing either two (low perceptual load) or four (high perceptual load) simultaneously presented lateralized sound stimuli. In separate task blocks, participants were instructed either to report whether the target was located on the left or the right side of the sound array (low spatial demand) or to indicate the exact target location (high spatial demand). Univariate alpha lateralization magnitude was neither affected by perceptual load nor by spatial demand. However, an analysis of onset latencies revealed that alpha lateralization emerged earlier in low (vs. high) perceptual load trials as well as in low (vs. high) spatial demand trials. Finally, we trained a classifier to decode the specific target location based on the multivariate alpha power scalp topography. A comparison of decoding accuracy in the low and high spatial demand conditions suggests that the amount of spatial information present in the scalp distribution of alpha-band power increases as the task demands a higher degree of spatial specificity. Altogether, the results offer new insights into how the dynamic adaption of alpha-band oscillations in response to changing task demands is associated with post-stimulus attentional processing.

## 1. Introduction

In everyday environments, containing multiple competing sensory inputs, focusing spatial attention on relevant information while ignoring or suppressing irrelevant information is crucial to engage in goal-directed behaviour. Consistently, covert shifts of spatial attention have been shown to improve various aspects of behavioural performance, including visual spatial acuity (reviewed by Anton-Erxleben & Carrasco, 2013), contrast sensitivity (Carrasco, Penpeci-Talgar, & Eckstein, 2000), or the rate of information accumulation (Carrasco & McElree, 2001). On the electrophysiological level, asymmetric modulations of parieto-occipital alpha-band power present a robust signature of spatial attentional orienting.

Typically, alpha-band power decreases contralateral to the attended location and / or increases over ipsilateral scalp sites. This phenomenon of alpha power lateralization has been found in response to anticipatory shifts of attention (Foxe, Simpson, & Ahlfors, 1998; Worden, Foxe, Wang, & Simpson, 2000), when retro-actively attending to working memory representations (Poch, Capilla, Hinojosa, & Campo, 2017; Schneider, Mertes, & Wascher, 2016), as well as during post-stimulus attentional processing (e.g., in auditory or visual search paradigms; Bacigalupo & Luck, 2019; Klatt, Getzmann, Wascher, & Schneider, 2018b).

Accumulating evidence suggests that scalp-level alpha-band activity not only reflects the attended hemifield but is tuned specifically to the attended visual field location (Bahramisharif, Heskes, Jensen, & van Gerven, 2011; Rihs, Michel, & Thut, 2007). Moreover, this spatial selectivity is also reflected in the retinotopic organization of alpha sources (Popov, Gips, Kastner, & Jensen, 2019). First evidence for comparable ‘spatial tuning’ of alpha-band oscillations in the auditory domain comes from a recent study by Deng and colleagues (Deng, Choi, & Shinn-Cunningham, 2020) who found that the topographic distribution of posterior alpha-band lateralization changes monotonically as the focus of auditory spatial attention shifts in space (see also Banerjee, Snyder, Molholm, & Foxe, 2011 for a comparison across sensory domains).

Notably, recent evidence suggests that the degree of spatial specificity reflected in the scalp distribution of alpha-band power also depends on the current task demands (Feldmann-Wüstefeld & Awh, 2019; Voytek et al., 2017). Specifically, compelling evidence comes from multivariate inverted encoding models (IEM), which can be used to quantify topographic patterns of alpha activity. Briefly, IEMs assume that the scalp distribution of (alpha-band) oscillatory power reflects the weighted sum of several spatially tuned neural populations (or spatial channels). By estimating the relative contribution of these channels (i.e., the channel weights), the model allows to eventually predict the response of the spatial channels from the distribution of alpha power across scalp electrodes. This results in a set of so-called “channel-tuning functions” that reflect the spatial selectivity of the underlying neuronal populations that contribute to the recorded scalp EEG (Foster, Sutterer, Serences, Vogel, & Awh, 2017). By means of IEMs, two studies of anticipatory visual attention demonstrated that the spatial selectivity of alpha activity increased when participants voluntarily focused on a narrow rather than a broad region of space (Feldmann-Wüstefeld & Awh, 2020) and scaled to the degree of certainty of a central cue that indicated the location of an upcoming target (Voytek et al. 2017).

Consistently, in an auditory spatial attention study, focusing on post-stimulus attentional processing, we found that task-demands shape the reliance on alpha-band mediated post-stimulus processing. That is, auditory post-stimulus alpha lateralization was only present in a spatially specific sound localization task, whereas it was absent in a simple sound detection paradigm (Klatt et al. 2018b, see also Deng et al. 2019). In the present study, we set out to further investigate to what extent attentional modulations of post-stimulus alpha power capture the spatial demands of a sound localization task on a more fine-grained scale. To this end, we varied both the perceptual load and the spatial demand of the task. That is, participants were asked to localize a target sound among a set of either two (low perceptual load) or four (high perceptual load) concurrently presented sounds in a lateralized sound array. In separate task blocks, they either indicated (a) whether the target was present on the left or the right side (i.e., two response options, low spatial demand) or (b) reported the exact target location (i.e., four response options, high spatial demand). On the behavioural level, we expected that high perceptual load (compared to low load) and high spatial demand (compared to low spatial demand) would present the more challenging listening situation, resulting in slower response times and lower sound localization accuracy. Beyond that, attempting to replicate previous results, we hypothesized that post-stimulus modulations of alpha-band power should index the attended target location, while the magnitude thereof should not be affected by perceptual load (Klatt et al., 2018b). This should be evident in a hemispheric lateralization of alpha-band power over parieto-occipital electrode sites in both low and high perceptual load trials.

Further, the critical aim of this study was to assess whether the required degree of behavioural spatial specificity (low vs. high spatial demand) affects the spatial specificity of the alpha power signal. If this is the case, this should be either evident in a modulation of alpha lateralization magnitude and / or captured by the scalp distribution of alpha-band power. Hence, we applied both univariate as well as multivariate analysis techniques to evaluate alpha-band power modulations depending on the spatial (and perceptual) demands of the task. Finally, we assessed alpha lateralization onset latencies to explore whether the time course of alpha-band activity is likewise modulated by the required degree of spatial specificity or perceptual load. Specifically, if slower sound localization performance in high spatial demand or high perceptual load conditions coincides with slower post-stimulus attentional processing, this should be reflected in delayed onset latencies of alpha lateralization. Such a time-resolved modulation of attentional alpha-band activity is, for instance, suggested by Foster and colleagues (Foster et al., 2017), who showed that the onset latency of location-selective alpha-band channel tuning functions (reconstructed from the topographic distribution of alpha-band oscillatory power) occurred later in time for trials with slow compared to fast responses as well as for a hard compared to an easier search condition.

## 2. Methods

### 2.1 Ethics statement

The study was approved by the Ethical Committee of the Leibniz Research Centre for Working Environment and Human Factors and conducted in accordance with the Declaration of Helsinki. All participants provided written informed consent prior to participation. The study procedure and analyses were not pre-registered prior to conducting the research. In the following sections, we report how we determined our sample size, all data exclusions, all inclusion/exclusion criteria, whether inclusion/exclusion criteria were established prior to data analysis, all manipulations, and all measures in the study.

### 2.2 Participants

19 participants were recruited to take part in the study. Hearing acuity was assessed using a pure-tone audiometry (Oscilla USB 330; Inmedico, Lystrup, Denmark), presenting eleven pure-tone frequencies in-between 125 Hz and 8000 Hz. One participant had to be excluded due to a unilateral, mild to moderate hearing impairment in the right ear (hearing thresholds of up to 35 – 50 dB hearing level). All other participants showed no signs of hearing impairment (hearing thresholds ≤ 25 dB). Another participant did not correctly follow the task instructions and was also excluded. Thus, the final sample included 17 subjects (mean age 23.29 years, age range 19-30, 9 female), all of which were right-handed as indicated by the Edinburgh Handedness Inventory (Oldfield, 1971). All participants had normal or corrected-to-normal vision, reported no history of or current neurological or psychiatric disorders and received course credit or financial compensation (10€/hour) for their participation. The above-mentioned inclusion and exclusion criteria were established prior to data collection. The sample size we aimed at was chosen to be comparable to previous publications from the lab that investigated similar electrophysiological measures (e.g., Klatt, Getzmann, Wascher, & Schneider, 2018b, 2018a). A sensitivity analysis, conducted using MorePower 6.0 (Campbell & Thompson, 2012), revealed that with a sample size of N=17, an alpha level of .05 and power of .8, the smallest effect size we can detect is η_p_^2^ = 0.36.

### 2.3 Experimental setup and stimuli

The experiment was conducted in a dimly illuminated, anechoic, and sound-attenuated room (5.0 × 3.3 × 2.4m^3^). Pyramid-shaped foam panels on ceiling and walls and a woolen carpet on the floor ensure a background noise level below 20dB(A). Participants were seated in a comfortable chair with their head position held constant by a chin rest. A semicircular array of nine loudspeakers (SC5.9; Visaton, Haan, Germany) was mounted in front of the subject at a distance of ~1.5 meters from the subject’s head and at a height of ~1.3 meters (approximately at ear level). Only five loudspeakers, located at azimuthal positions of −90°, −30°, 0°, 30°, and 90° respectively, were used for the present experimental setup. A red, light-emitting diode (diameter 3 mm) was attached right below the central loudspeaker. The diode remained turned off during the experiment but served as a central fixation target.

As sound stimuli, eight familiar animal vocalizations (‘birds chirping’, ‘dog barking’, frog croaking’, ‘sheep baaing’, ‘cat meowing’, ‘duck quacking’, ‘cow mooing’, ‘rooster crowing’) were chosen from an online sound archive (Marcell, Borella, Greene, Kerr, & Rogers, 2000). The original sounds were cut to a constant duration of 600 ms (10 ms on/off ramp), while leaving the spectro-temporal characteristics unchanged. The overall sound pressure level of the sound arrays, containing either two or four concurrently present sounds, was about 63 dB(A) and 66 dB(A), respectively. The target sounds, presented in isolation from a central position, had a sound pressure level of 60 dB(A).

### 2.4 Procedure, task, and experimental design

The experiment consisted of an auditory search paradigm implementing a sound localization task. The sequence of events in a given trial is depicted in Figure 1. Each trial began with a silent period of 500 ms. Then a sound stimulus (i.e., a target cue) was presented from a central position (0° azimuth angle) for 600 ms, indicating which animal vocalization will serve as a relevant target sound in a given trial. The latter was followed by a 1000 ms silent inter-stimulus-interval and a sound array (600 ms). The sound array contained either two (i.e., *low perceptual load*, 50%) or four (i.e., *high perceptual load*, 50%) simultaneously present lateralized sound stimuli. Responses were permitted immediately after sound array onset and until the end of a 1600 ms response interval. The latter was followed by another 300 ms silent interval. In total, each trial lasted for 4600 ms.

**Figure 1.**
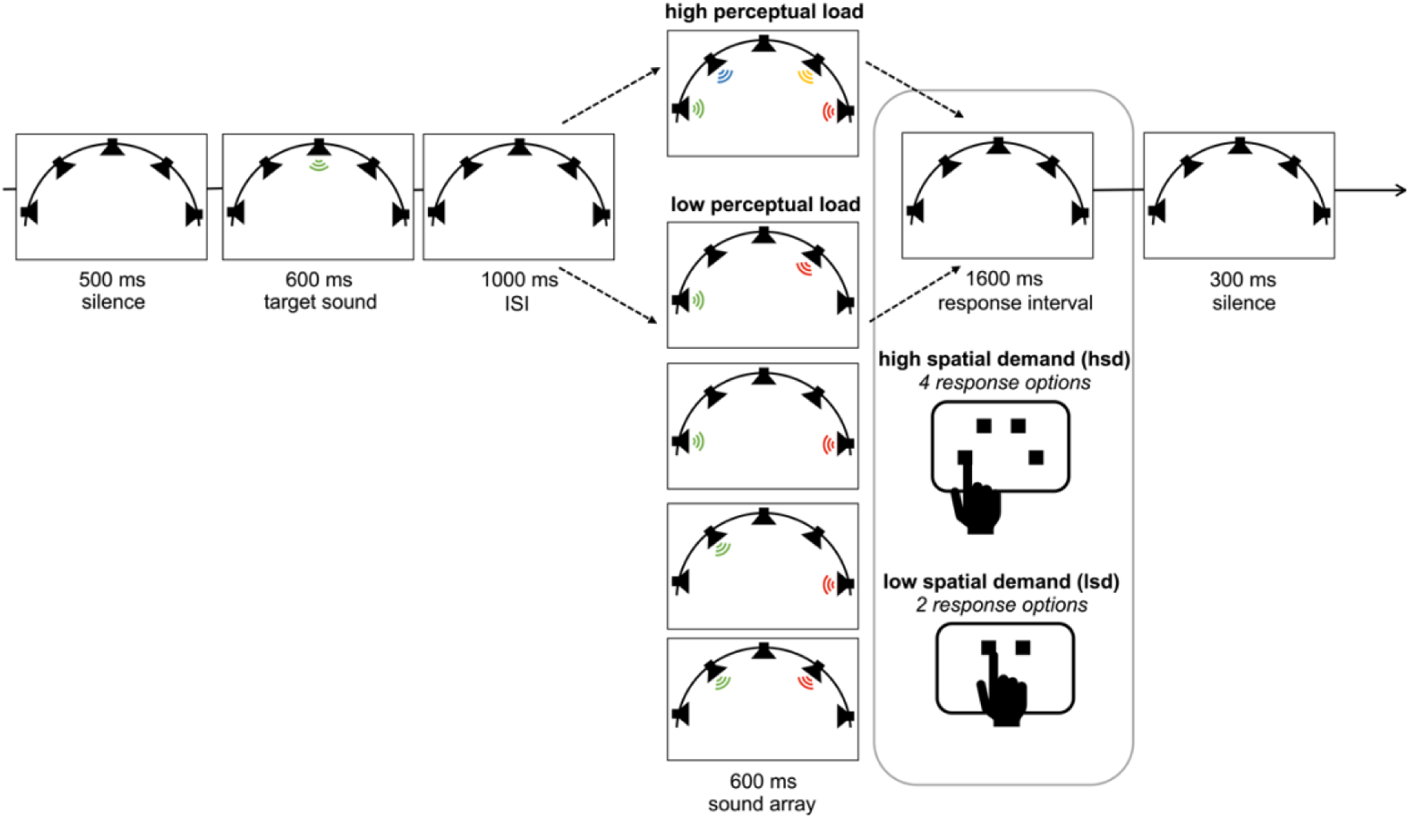
Schematic illustration of the experimental design. A centrally presented target cue indicated the relevant target in a given trial. Then, a sound array appeared, containing either two or four simultaneously present sounds from lateralized positions. In different task blocks, participants were asked to either indicate whether the target was presented on the left or the right side (low spatial demand) or to report the exact target location (high spatial demand). Responses were permitted immediately following sound array onset (i.e., there was no need to withhold the response until the onset of the response interval). In both task blocks, it was also possible that the sound array did not contain the target (i.e., target-absent trial). In this case, participants withheld their response. ISI = inter-stimulus-interval.

In low perceptual load trials, the two sounds could occur at either of the four lateralized loudspeaker positions (−90°, −30°, 30°, 90° azimuth), with the restriction that the two sounds (i.e., the target and a non-target sound) were always present in different hemi-fields. Accordingly, in high perceptual load trials all four lateralized active loudspeakers (−90°, −30°, 30°, 90° azimuth) were used. Depending on the task condition, participants received slightly different task instructions: In the *low spatial demand (lsd)* condition, participants were instructed to indicate whether the target sound was present on the left versus right side (i.e., two response options: left vs. right) or to withhold their response if the target sound was not present (i.e., target-absent trials). In the *high spatial demand (hsd)* condition, participants were asked to indicate the exact target location (i.e., four response options: inner-left, outer-left, inner-right, outer-right) or to withhold their response if the target sound was not present. Target-absent trials were included to ensure that selectively listening to the input from only one side of the stimulus array (i.e., left or right) presented no viable strategy in low spatial demand task blocks. Specifically, if the sound array always contained a target sound in low spatial demand blocks, subjects could be inclined to simply infer that the target was located on the left side solely because they didn’t perceive it on the right side (or vice versa).

Participants indicated their response by pressing one out of four buttons, arranged in a semi-circular array on a response pad. In the high-spatial demand condition, each button corresponded to one of the loudspeaker positions, such that participants had to press the left-most button when the target was presented at the left-most loudspeaker and so on. In low spatial demand trials, participants only used the two inner response buttons (i.e., the left button for left-target responses, the right button for right-target responses). Participants were instructed to always respond as accurately and as fast as possible, using the index finger of their right hand. To minimize horizontal eye movements during the EEG-recording, participants were instructed to fixate a centrally positioned LED.

Each of the spatial demand conditions (i.e., low vs. high spatial demand) consisted of 672 trials, containing both low (50%) and high (50%) perceptual load trials in randomized order. Short, self-paced breaks after every 224 trials and in-between conditions were conducted to prevent fatigue. The order of conditions was counterbalanced across participants, such that n = 8 subjects first completed the low-spatial demand condition and n = 9 subjects first completed the high-spatial demand condition. Prior to the beginning of each condition participants completed 40 practice trials to familiarize with the task. All participants were presented with the same semi-randomized selection of trials. Critically, in both spatial demand conditions the same selection of 672 trials was presented, but in a different, randomized order. This assured that all differences between conditions could be ascribed to the task manipulations rather than differences in the stimulus materials. Each of the eight animal vocalizations served as the target equally often (i.e., 84 times per condition). In addition, the target sound appeared equally often at each of the four possible sound speaker locations (i.e., 56 times per location and perceptual load per condition). This also ensured that the number of left (1/3) vs. right (1/3) responses in low-spatial demand trials as well as the number of outer-left (1/5), inner-left (1/5), inner-right (1/5), and outer-right (1/5) responses in high-spatial demand trials was counterbalanced across subjects. Target-absent trials constituted 1/3^rd^ and 1/5^th^ of all trials in low and high spatial demand task blocks, respectively. The timing of the stimuli was controlled by custom-written software. Participants did not receive feedback during the experiment.

Taken together, the present study comprised a 2 x 2 repeated-measures design, including the within-subject factors *spatial demand* (low vs. high spatial demand) and *perceptual load* (low vs. high perceptual load). Note that there are different ways of defining perceptual load (for a review see Murphy, Spence, & Dalton, 2017). Here, we refer to *perceptual load* as the number of items in the search display.

### 2.5 EEG data acquisition

The continuous EEG data were recorded from 58 Ag/AgCl passive scalp electrodes (ECI Electrocap, GVB-geliMED GmbH, Bad Segeberg, Germany) as well as from left and right mastoids. Electrode positions corresponded to the international 10-10 system. The electrooculogram (EOG) was simultaneously recorded from four additional electrodes, placed lateral to the outer canthus of each eye as well as below and above the right eye. The ground electrode was placed on the center of the forehead, right above the nasion. The average of all electrodes served as the online-reference. The data were recorded using a QuickAmp-72 amplifier (Brain products, Gilching, Germany) and digitized at a sampling rate of 1 kHz. During the preparation of the EEG cap, all electrode impedances were kept below 10 kΩ.

### 2.6 Data analysis

If not stated otherwise, all data analyses were performed using custom MATLAB (R2021b) code and built-in functions from the *Statistics and Machine Learning Toolbox.* In a few specific cases, R (v3.6.1) and RStudio (2021.09.2) were used (see references to specific R packages below). The significance of all tests was evaluated at an alpha level of .05. Because the *F*-distribution is always asymmetric, reported *p*-values associated with repeated-measures analysis of variance (ANOVA) are directional (Winter, 2011). Partial Eta Squared (η_p_^2^) and Hedges’ g (denotes as *g*, Hentschke & Stüttgen, 2011) are provided as standardized measures of effect size for ANOVAs and follow-up paired sample *t*-tests.

#### 2.6.1 Behavioral

The behavioral parameters that were analyzed were response times and accuracy (i.e., percentage of correct responses). Please note that target-absent trials did not require a button press because subjects were instructed to withhold their response. Hence, we cannot safely distinguish whether a non-response in a target-absent trial was due to the correct perception that no target was presented *or* due to occasional attentional lapses, causing the subject to “miss” a trial. Consequently, to compute the percentage of correct responses, we only considered target-present trial, which always required a button press. To compute mean response times, only correct target-present trials were considered. Pre-mature responses (< 200 ms post-sound array onset) were excluded from further analyses. Mean RTs and accuracy measures per subject and condition were submitted to a repeated-measures ANOVA. Spatial demand and perceptual load served as within-subject factors.

#### 2.6.2 EEG

All EEG data processing was performed using the open-source toolbox EEGLAB (v14.1.2; Delorme & Makeig, 2004) in combination with custom MATLAB (R2021b) code.

##### 2.6.2.1 Preprocessing

Initially, continuous segments of −1 to +1 seconds surrounding boundary events as well as the DC offset were removed from the data. Then, the continuous EEG data were band-pass filtered, using a non-causal, high-pass and a low-pass Hamming windowed sinc FIR filter (*pop_eegfiltnew* function). The lower edge of the frequency pass band was set to 0.1 Hz (filter order: 33000, transition band-width: 0.1 Hz, −6dB cutoff: 0.05 Hz) and the higher edge of the frequency pass band to 30 Hz (filter order: 440, transition band-width: 7.5 Hz, −6dB cut-off: 33.75 Hz). Early-stage preprocessing was then performed using the PREP pipeline (Bigdely-Shamlo, Mullen, Kothe, Su, & Robbins, 2015), which essentially consists of three steps: it performs an initial clean-up, determines and removes a robust reference signal, and interpolates bad channels with a low signal to noise ratio. For an extensive documentation of the single steps, please see Bigdely-Shamlo et al. (2015). Only scalp EEG channels were used for evaluation of noisy channels and for computation of the robust reference, while all channels (including mastoids and EOG channels) were re-referenced. On average, 3.8 channels (SD = 2.3) were identified as bad and interpolated prior to subtracting the computed “true” reference. This includes a total of three channels (across two subjects) that were manually interpolated prior to running the PREP algorithm, because the latter did not identify the respective channels as flat channels. For channel interpolation, the PREP pipeline applies spherical spline interpolation as implemented in the *eeg_interp()* function (Perrin, Pernier, Bertrand, & Echallier, 1989), which is based on the electrode location coordinates of the international 10-10 system. The same algorithm was used to manually interpolate the three channels that were not identified as flat channels by the PREP algorithm. A total of three channels (in two subjects) belonging to the posterior electrode cluster of interest that was used for statistical analysis (see section 2.6.2.3) were marked as bad and thus, interpolated during this procedure.

For artifact rejection, an independent component analysis (ICA) was run on the dimensionality reduced data (using a basic PCA implementation). To speed up and improve ICA decomposition, the continuous data were down-sampled to 200 Hz and high-pass filtered at 1 Hz (Winkler, Debener, Muller, & Tangermann, 2015), using a non-causal Hamming windowed sinc FIR filter (filter order: 3300, transition band-width: 1 Hz, −6dB cutoff: 0.5 Hz) prior to running the ICA algorithm. Then, data epochs were extracted, ranging from −1000 to 5000 ms relative to target cue onset. In addition, major artefacts and extremely large potential fluctuations were removed before running ICA, using the automatic trial-rejection procedure implemented in EEGLAB (i.e., function *pop_autorej*). The latter rejects data epochs, containing data values exceeding a given standard deviation threshold by means of an iterative procedure (probability threshold: 5 SD, maximum proportion of total trials rejection per iteration: 5%, threshold limit: 500 µV). Because interpolating channels prior to ICA introduces rank-deficiency, the number of to-be extracted ICs was manually reduced by the number of interpolated channels + 1 (to account for the dependency introduced by the average reference). To identify artefactual independent components (ICs), the EEGLAB plug-in ICLabel (v1.1, Pion-Tonachini, Kreutz-Delgado, & Makeig, 2019), was applied. ICLabel assigns a label vector to each IC, indicating the probability that an IC belongs to any of seven possible categories: brain, muscle, eye, heart, line noise, channel noise, or other. All ICs that received a probability estimate below 50% for the brain category were marked as “artefactual” (Arnau et al., 2020; Reiser, Wascher, Rinkenauer, & Arnau, 2020). Further, one restriction applied: ICs with the maximum probability estimate for the category brain were not rejected even if the probability was below 50% to avoid that meaningful brain activity was removed from the signal. The obtained ICA decomposition and the ICLabel results were copied to the original, continuous dataset (band-pass filtered and re-referenced) with a 1 kHz sampling rate. The latter was segmented into epochs ranging from −1000 to 5000 ms relative to target cue onset and baseline-corrected, using the pre-stimulus period of −200 to 0. Then, ICs that were previously identified as artefactual (On average, 32.82 ICs; SD = 4.41) were removed from the signal. Finally, to remove remaining artefacts that were not accounted for by the ICA-based artefact rejection procedure, trials with large fluctuations (−150 / + 150 µV) were excluded, using the automated EEGLAB algorithm *pop_eegthresh()*. Moreover, trials with premature responses (i.e., response time < 200 ms) were excluded (on average 5 trials, SD = 9). Not considering target-absent trials (i.e., irrelevant for the present analyses), on average, 216 (lsd-low, SD = 11), 216 (lsd-high, SD = 9), 213 (hsd-low, SD = 10), and 213 (hsd-high, SD = 11) *target-present-trials* passed the artefact correction per subject. Specifically, 211 (lsd-low, SD = 14), 190 (lsd-high, SD = 18), 205 (hsd-low, SD = 13), and 176 (hsd-high, SD = 21) of those target-present trials were correct trials, and thus entered the univariate EEG analysis. This corresponds to, on average, 106 (lsd-low, SD = 7), 95 (lsd-high, SD = 9), 103 (hsd-low, SD = 7), and 88 (hsd-high, SD = 10) trials per target hemifield.

##### 2.6.2.2 Time-frequency decomposition

The time-frequency decomposition of the processed EEG data was computed using Morlet wavelet convolution as implemented in the build-in EEGLAB STUDY functions (i.e., *newtimef.m*). Specifically, the segmented EEG signal was convolved with a series of complex Morlet wavelets. The frequencies of the wavelets ranged from 4 Hz to 30 Hz, increasing logarithmically in 52 steps. A complex Morlet wavelet is defined as a complex sine wave that is tapered by a Gaussian. The number of cycles, that defines the width of the tapering Gaussian, increased linearly as a function of frequency by a factor of 0.5. This procedure accounts for the trade-off between temporal and frequency precisions as a function of the frequency of the wavelet. The number of cycles at the lowest frequency was 3; the number of cycles at the highest frequency was 11.25. The time period in-between −400 and −100 ms relative to target cue onset served as a spectral baseline.

##### 2.6.2.3 Alpha power lateralization

Spatial shifts of attention following the onset of the sound array were quantified by assessing lateralized modulations of posterior alpha-band power (8-12 Hz). Specifically, the difference between contralateral and ipsilateral alpha power at a cluster of posterior electrodes, comprising PO7/8, P7/8, P3/4, and PO3/4, was calculated separately for each condition and each subject. The selection of electrodes was based on previous studies of post-stimulus, posterior alpha lateralization (Klatt, Getzmann, Begau, & Schneider, 2019; Schneider, Göddertz, Haase, Hickey, & Wascher, 2019), except that P5/P6 were not part of the present electrode setup and thus, electrodes P3/4 were included in the electrode cluster instead. Given that post-stimulus alpha power asymmetries have been shown to appear as a relatively long-lasting, sustained effect (Klatt et al., 2018a), the mean contralateral-minus-ipsilateral differences in power were extracted in a broad 400 ms-time window, ranging from 546 to 961 ms following sound array onset.^1^ The time window was set around the peak in the grand average contralateral minus ipsilateral difference waveform across all conditions and subjects. The peak was defined as the point in time at which the difference waveform (following sound array onset, 1600 ms – 3000 ms) reached its most negative amplitude value. The resulting analysis time window is consistent with our earlier work (Klatt et al., 2018a). Notably, although this approach to determine the analysis time window is data-driven, the comparisons between conditions remain unbiased (Luck & Gaspelin, 2017). The mean power values per subject and condition were then submitted to a repeated-measures ANOVA, including the within-subject factors *spatial demand* and *perceptual load* to assess their effect on alpha lateralization magnitude.

##### 2.6.2.4 Alpha lateralization onset latencies

To quantify the time course of alpha lateralization, we used a combination of the fractional area technique (Kiesel, Miller, Jolicœur, & Brisson, 2008; Luck, 2014) and a jackknife approach (Luck, 2014; Miller, Patterson, & Ulrich, 1998). That is, for each condition, *n* subaverage contralateral minus ipsilateral difference waveforms were created, using a subsample of *n-1* waveforms (i.e., each participant was omitted once). In each of these subaverage waveforms, the point in time at which the negative area under the curve reached 20% and 50%, respectively (i.e., Fractional Area Latency, denoted as FAL) was measured, using the MATLAB function *latency.m* by Liesefeld (2018). Negative area was measured relative to zero and in-between a broad time window from 1600 to 3000 ms post-cue-onset (i.e., 1600 ms corresponds to sound array onset). Note that reported mean latency differences (denoted as *D*) correspond to the differences in onset latency between conditions, measured in the condition-grand averages. According to Miller, Patterson, & Ulrich (1998), the jackknife-based *SE_D_* was calculated as follows:

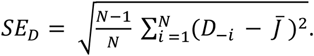

*D_−i_* (for I = 1, …, N, with N representing the sample size) denotes the latency difference for the subsample, including all subjects except for subject *i*. 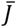 is the mean difference across all subsamples (i.e., 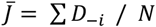).

The 20%-FAL and 50%-FAL values were submitted to separate repeated-measures ANOVAs, including the within-subject factors *spatial demand* and *perceptual load.* Because the use of subsample average measures artificially reduces the error variance, the error terms in the respective ANOVA will be underestimated, while the F-values will be overestimated. To account for this bias, the *F*-correction according to Kiesel, Miller, Jolicœur, & Brisson (2008) was applied. Corrected *F*-values are denoted as F_corr_. The corresponding *p*-value for the corrected *F* statistic was computed using the online calculator by Soper (2020). In accordance with Richardson (2011), partial eta squared was computed based on the corrected F-values and the degrees of freedom (cf. formula 8 in Richardson, 2011).

Please note that the main aim of the present analysis of onset latency measures was to assess differences in time course of alpha lateralization between the experimental conditions. However, the estimated latency measures should not be interpreted as reflecting the true onset time of the underlying attentional process. This precaution applies for two reasons: First, the temporal resolution of event-related spectral perturbations is considerably lower compared to standard ERP analysis. Second, non-causal filters, as applied here to the continuous raw EEG data, have been shown to affect the onset latency of time-series data considerably (VanRullen, 2011, but see also Rousselet, 2012). Critically, as the filter should affect all conditions to the same extent, the differences between conditions can still be reliably interpreted.

##### 2.6.2.5 Non-lateralized, posterior alpha power desynchronization

Event-related desynchronization (ERD) of alpha-band activity resulting in low levels of alpha power has been linked with states of high excitability and thus, is thought to reflect functional engagement and information processing (see e.g., Fukuda, Mance, & Vogel, 2015; Krause et al., 2000; Hanslmayr, Spitzer, & Bäuml, 2009). Hence, in the present study, posterior alpha ERD served as a measure of cognitive task demands. Mean alpha ERD amplitude per condition and subject was measured in-between 1698 ms and 2698 ms relative to target cue onset (i.e., 98 – 1098 ms relative to sound array onset) at electrode Pz. The time window that served as the basis for the statistical analysis was determined using a collapsed localizer approach (Luck & Gaspelin, 2017). That is, we assessed the negative peak in the grand average waveform across conditions in a broad time-window from 1600 ms to 3000 ms (relative to target cue onset, i.e., the same time window used to measure the area under the curve for fractional area latency measurement). In accordance with previous studies which characterized post-stimulus alpha ERD as a broad, sustained response (Fukuda, Mance, & Vogel, 2015; Krause et al., 2000; Hanslmayr, Spitzer, & Bäuml, 2009), a 1000 ms time window (i.e., +/− 500 ms) around the resulting peak latency of 2194 ms (i.e., 594 ms following sound array onset) constituted the measurement time window. Mean alpha power values were then submitted to a repeated-measures ANOVA, including the within-subject factors *spatial demand* (high vs. load) and *perceptual load* (high vs. low).

##### 2.6.2.6 Decoding analysis

We attempted to decode the exact location (i.e., outer-left, inner-left, inner-right, outer-right) of the target sound based on the scalp distribution of alpha-band EEG power. The decoding procedure was applied separately for the *low* vs. the *high spatial demand* condition to investigate whether the ‘amount’ of spatial information reflected in the scalp topography of alpha-band power is modulated by the spatial demands of the task. The factor *perceptual load* was not considered in the decoding analysis for two reasons: First, the decoding analysis primarily aimed to follow up on similar analyses that investigated anticipatory (i.e., pre-stimulus) modulations of alpha power (Feldmann-Wüstefeld & Awh, 2019; Voytek et al., 2017) which provided evidence that the scalp distribution of alpha-band power is modulated by the spatial specificity of the task demands. In addition, performing separate decoding analyses for the different perceptual load conditions would have resulted in a substantially reduced number of trials available for each decoding routine. Further, only correct trials were considered for the decoding analysis. This ensures that differences in accuracy between the two spatial demand conditions to not affect the results.

The applied classification routine was adapted from Bae and Luck (2018). First, to isolate the alpha-band signal of interest, the segmented EEG at all scalp electrodes was bandpass filtered at 8 to 12 Hz, using EEGLAB’s *eegfilt()* function, which applies two-way least-squares finite impulse response (FIR) filtering. Then, we submitted the bandpass filtered EEG data to a Hilbert transform to obtain the magnitude of the complex analytic signal. The latter was squared to compute the total power in the alpha frequency band (i.e., 8-12 Hz) at each time point. ^2^ Subsequently, to increase the efficiency of the analysis and decrease computation time, the data was subsampled, keeping only every 20^th^ data point in-between −500 and 4500 ms relative to target sound onset (i.e., corresponding to a sampling rate of 50 Hz). This results in a 4-dimensional data matrix for each participant, including the dimensions of time (250 time points), location (4 different categories), trial (varies depending on the subject, in-between 64 and 110 trials for each location), and electrode site (the 57 scalp channels).

To classify the location of the target sound based on the scalp topography of the alpha power signal over the 57 scalp electrodes (mastoids and EOG electrodes were excluded), we used a combination of a support vector machine (SVM) and error-correcting output codes (ECOC; Dietterich & Balkiri, 1995). The ECOC model, implemented using the MATLAB function *fitcecoc()*, combines the results from multiple binary classifiers and thus, solves multiclass categorization problems. Classifications were performed within subjects and using trial averages rather than single-trial data. The latter increases the signal-to-noise ratio in the classifier input and has been shown to result in higher (although more variable) decoding accuracies (Adam, Vogel, & Awh, 2020).

Specifically, decoding was performed separately for each of the 250 time points in-between −500 and 4500 ms relative to target sound onset. At each time point, 50 iterations of the classification analysis were performed; on each iteration, the data were sorted into four ‘location bins’, containing only trials with the same target location. In each location bin, the trials were randomly divided into three equally sized sets of trials. That is, to ensure that an equal number of trials was assigned to each of the three sets for each location bin, the minimum number of trials per subject for a given location bin was determined (denoted as *n*), and *n* / 3 trials were assigned to each set. In case the total trial number for a given location was not evenly divisible by three, excess trials were randomly omitted. The trials for a given location bin were averaged, resulting in a matrix of 3 (subsample averages) x 4 (location bins) x 57 (electrodes) to be analyzed for each time point. Two of the three subsample averages (for each target location) served as the training set, while the remaining subsample average (for each target location) was assigned to the testing dataset. In the training phase, the data from the two (of the total three) subsample averages was simultaneously submitted to the ECOC model with known location labels to train four SVMs (one for each location). Each SVM was trained to perform a binary classification, that is, to discriminate one specific location from all other locations. Subsequently, in the test phase the unused data (i.e., the subsample averages that were reserved for testing) was fed into the set of four trained SVMs to classify which of the 4 locations served as the target location in each of the subsample averages; hence, by combing the results from multiple binary classifiers, a multi-class categorization problem is solved. Specifically, the MATLAB *predict()* function was used to classify the input data by minimizing the average binary loss across the four trained SVMs. Essentially, the output of the *predict()* function provides a location label for each of the four remaining subsample averages in the testing dataset. By comparing the true location labels to the predicted location labels, decoding accuracy was computed.

Decoding was considered correct if the classifier correctly determined which one of the four possible locations was the target location. Thus, chance level decoding accuracy was at 25%. This training-and-testing process was applied three times such that each subsample average served as the testing dataset once. Finally, decoding accuracy was collapsed across the four locations, the three cycles of cross-validation, and the 50 iterations, resulting in a decoding percentage for each time point. After obtaining a decoding percentage for all time points of interest, a five-point moving average was applied to smooth the averaged decoding accuracy values and to minimize noise.

##### 2.6.2.7 Statistical analysis of decoding accuracy

Although decoding was performed for all time points in-between −500 to 4500 ms relative to sound onset, the statistical analysis focused on the time interval following sound array presentation until the end of the maximal response interval (i.e., 1600 – 3800 ms relative to sound onset). We restricted the statistical analysis to this time interval because the goal was to test decoding accuracy during the post-stimulus interval (i.e., when post-stimulus attentional processing takes place). In addition, because participants did not have any knowledge about *where* the target is going to appear prior to sound array onset, there should be no location-specific information present in-between target cue and sound array-onset. Briefly, the statistical analysis of decoding accuracy comprised two separate approaches: First, to confirm that the scalp topography of post-stimulus alpha-band power contains information about the target location, we compared decoding accuracy to chance level (i.e., 25% – because we used 4 locations) at each time point. This was done separately for the two spatial demand conditions. Second, we compared decoding accuracy in the low and high spatial demand condition to evaluate whether the amount of spatial information that is reflected in the scalp topography of alpha-band power is sensitive to the spatial demands of the task. At both stages, we controlled for multiple comparisons (see below for details).

##### 2.6.2.8 Decoding accuracy within conditions

We used a non-parametric cluster-based permutation analysis to compare decoding accuracy to chance level (i.e., 25%) at each time point. Here, we adopted the corrected analysis code provided by Bae and Luck (2019), accounting for the presence of autocorrelated noise in the data. Using one-sided one sample *t*-tests, the average decoding accuracy across subjects was compared to chance level, separately for each time-point.

Because SVM decoding does not produce meaningful below-chance decoding results, a one-sided *t*-test is justified. Then, clusters of at least two adjacent time points with a significant single-point *t*-test (i.e. *p* < .05) were identified. The *t*-values within a given cluster were summed, constituting the so-called cluster mass. To determine whether a given cluster mass is greater than what can be expected under the null hypothesis, we constructed a null distribution of cluster-level t-mass values using permutation tests. Critically, to reduce computation time, we randomly permuted the target labels at the stage of testing the decoding output, rather than prior to training the classifier. Specifically, from an array containing all possible target labels (1, 2, 3, 4), we randomly sampled an integer as the simulated response of the classifier for a given target location. If the response matched the true target value, the response was considered correct. This yields an estimate of the decoding accuracy values that would by obtained by chance if the decoder randomly guessed the target location. Critically, to reflect the temporal autocorrelation of the continuous EEG data, the same randomly sampled target position label was used for all time points in a given subaverage. Overall, this sampling procedure was repeated 600 times (4 locations x 3 cross-validations x 50 iterations) and for each time point of interest in-between 1600 ms to 3800 ms. The scores for each time point were averaged to obtain the mean simulated decoding accuracy, resulting in a time series of decoding accuracy values.

Analogous to the procedure that was applied to the actual EEG data, the latter was smoothed using a five-point running average filter. The procedure was repeated 17 times, to obtain a simulated decoding accuracy time series for each of our 17 participants. Then, using the simulated decoding accuracy time series, the maximum cluster mass was computed, using the procedure described above. That is, if there was more than one cluster of significant *t*-values, the mass of the largest cluster was selected.

Finally, this procedure (i.e., simulating decoding accuracy that would be obtained by chance) was iterated 10,000 times to produce a null distribution of cluster mass values. For each cluster in the decoding results, the obtained cluster *t* mass was compared to the distribution of cluster t mass values that was constructed under the assumption that the null hypothesis is true. If the observed cluster *t* mass value was larger than the 95^th^ quantile of the null distribution (i.e., α = .05, one-tailed), the null hypothesis was rejected and decoding accuracy was considered above chance. Note that this procedure was separately applied to both the low spatial demand condition and the high spatial demand condition.

To find the *p*-value associated with a specific cluster, we examined where within the null distribution does each observed cluster *t* mass value fall. That is, the *p*-value was based on the inverse percentile (computed using the *invprctile*() function) of the observed cluster-level *t* mass within the null distribution. If the observed cluster-level *t*-mass value exceeded the maximum cluster-level *t*-mass of the simulated null distribution, the respective *p*-value is reported as *p* < 10^−4^. The latter corresponds to the resolution of the null distribution (i.e., 1 / number of permutations).

##### 2.6.2.9 Decoding Accuracy in low versus high spatial demand blocks

To investigate, whether or not the amount of spatial information reflected by the scalp topography of alpha power differs depending on the spatial demands of the task, decoding accuracy in the two task conditions was compared, using a cluster-corrected sign-permutation test. To this end, the *cluster_test()* and *cluster_test_helper()* functions provided by Wolff, Jochim, Akyürek, and Stokes (2017) were applied. The sign-permutation test is a non-parametric test that makes no assumption of the distribution of the data. As input data, the same time window that was also used for the statistical analysis of decoding accuracy within conditions was selected (i.e., 1600 – 3800 ms). Specifically, the *cluster_test_helper()* function generates a null distribution by randomly flipping the sign of the input data of each participant with a probability of 50%. This procedure was repeated 10,000 times. The resulting distribution served as input to the *cluster_test()* function, identifying those clusters in the actual data that are greater than would we expected under the null hypothesis. The cluster-forming threshold as well as the cluster significance threshold were set to p < .05. Because we had a clear hypothesis regarding the direction of the effect (that is, decoding accuracy in the high spatial demand condition should be higher compared to the low spatial demand condition), the cluster-corrected sign-permutation test was one-sided.

In addition, to assess the overall difference in decoding ability within the post-stimulus period, the decoding accuracy was averaged across time in the approximate time window that resulted in significant within-condition decoding results across the two large clusters in both conditions (i.e., 1920 – 3380 ms) and submitted to a one-sided permutation test. To this end, the *GroupPermTest()* function provided by Wolff et al. (2017) was applied (using nSims = 10,000 permutations).

##### 2.6.2.10 Response-locked analyses

Several observations in the univariate and the multivariate analysis of alpha power prompted us to add the following post-hoc analyses^3^:

First, we time-locked the processed EEG data to the response, creating new epochs ranging from −2800 to 800 ms relative to the response. Then, Morlet wavelet convolution was applied as described in section 2.5.2. The resulting alpha-band event-related spectral perturbations (ERSPs) ranged from −2382 to 382 ms. Analogous to the stimulus-locked analysis, mean alpha-band power was computed in a broad 400 ms time-window in-between −182 to 224 ms^4^ (relative to the response). The time window was set around the peak in the grand average contralateral minus ipsilateral difference waveform across all conditions at 21 ms. The peak was determined by identifying the point in time at which the difference waveform (in-between −500 to 380 ms) reached its most negative amplitude value. Mean alpha power values for each condition were submitted to a repeated-measures ANOVA, including the within-subject factors spatial demand and perceptual load. Further, we obtained estimates of 20%- and 50%-fractional area latency, using a jackknifing approach (Miller et al., 1998), as described above. Specifically, the point in time at which the negative area in-between −500 to 380 ms reached 20% and 50%, respectively, was determined. The latency estimates were submitted to a repeated-measures ANOVA, including the within-subject factors spatial demand and perceptual load.

Finally, we applied the same decoding procedure as describe above, using the response-locked alpha-band ERSPs as classifier input. Again, a non-parametric cluster-based permutation analysis was applied to compare decoding accuracy to chance level. Considering the time-course of the univariate, response-locked alpha-band ERSPs, only time points in-between −500 and 380 ms (i.e., last sampling point in the ERSP data) were included in the statistical analysis. To contrast decoding accuracy between the two spatial demand conditions, we obtained the average decoding accuracy in a broad time window surrounding the response (−340 to 360 ms, i.e., the time-period that resulted in significant within-condition decoding across both conditions) and performed a one-sided permutation test (as described above).

### 2.7 Data/code availability statement

Stimuli and code for this study can be found at https://osf.io/a8f6y/. Data will be publicly stored in a Zenodo repository. Access will be granted by the corresponding author upon signing a data user agreement.

## 3 Results

### 3.1 Behavioral data

Behavioral results are displayed in Figure 2. The analysis of response times revealed a main effect spatial demand, *F*(1,16) = 72.49, *p* < .001, η_p_^2^ = 0.82, with slower responses in high spatial demand blocks (M = 837.67 ms, SD = 95.29) compared to low spatial demand blocks (M = 837.67 ms, SD = 126.83). In addition, there was a significant main effect of perceptual load, *F*(1,16) = 158.36, *p* < .001, η_p_^2^ = 0.91, with slower responses in high-load trials (M = 848.38 ms, SD = 119.32) compared to low-load trials (M = 707.56 ms, SD = 101.61). For response times, there was no significant interaction of spatial demand and perceptual load, *F*(1,16) = 3.19, *p* = .093, η_p_^2^ = 0.17. A nearly analogous pattern of results was revealed by the analysis of the percentage of correct responses. That is, participants responded more accurately in low spatial demand blocks (M = 92.38 %, SD = 4.82) compared to high spatial demand blocks (M = 88.75 %, SD = 4.83), *F*(1,16) = 20.97, *p* < .001, η_p_^2^ = 0.57). In addition, the percentage of correct responses was higher in low-load trials (M = 96.45 %, SD = 2.87), compared to high-load trials (M = 84.68 %, SD = 7.40), *F*(1,16) = 54.19, *p* < .001, η_p_^2^ = 0.77. Further, a significant interaction of spatial demand and perceptual load, *F*(1,16) = 11.11, *p* = .004, η_p_^2^ = 41, complements the descriptive observation that the difference in accuracy between low and high perceptual load was slightly greater in high spatial demand blocks (M = 13.63 %, SD = 7.12) than in low spatial demand blocks (M = 9.93 %, SD = 6.86).

**Figure 2.**
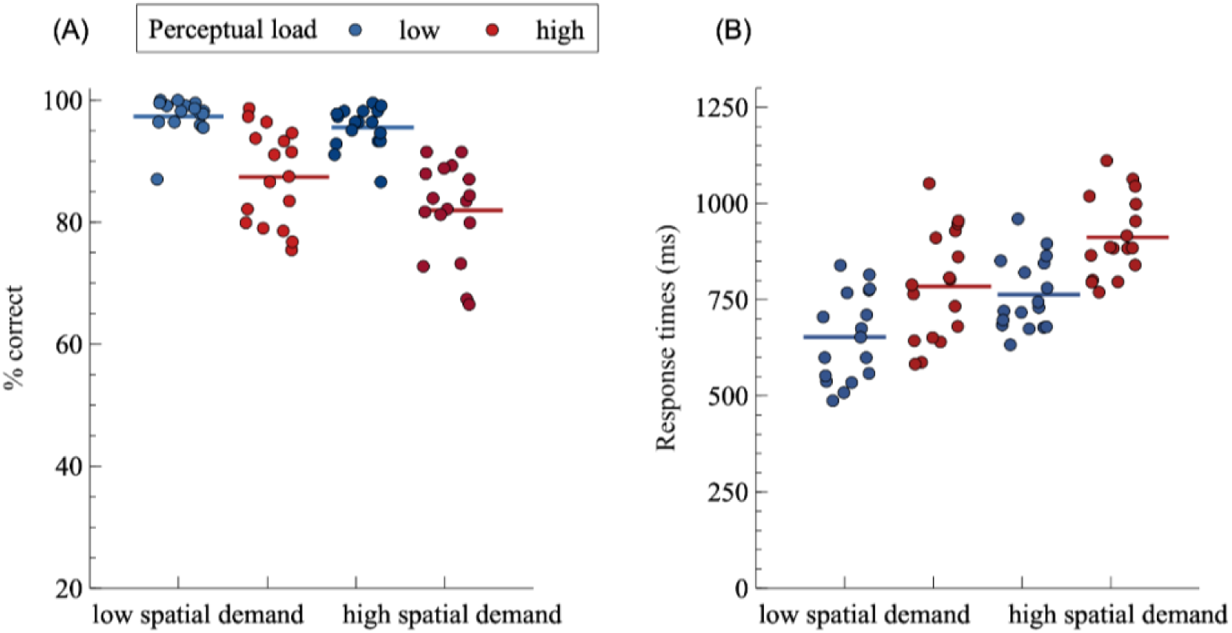
Behavioral performance. Solid, horizontal lines indicate the mean percentage of correct responses (A) or mean response times (B) in a given condition. Colored dots correspond to individual response measures. Please note that the y-axis for the % of correct responses does not originate at 0.

### 3.2 Alpha power lateralization

Figure 3A illustrates the time course of the contralateral minus ipsilateral differences in alpha power at a cluster of posterior scalp electrodes. A repeated-measures analysis of the mean alpha power amplitudes in-between 546 to 961 ms post-sound array onset revealed no significant modulation by spatial demand, *F*(1,16) = 1.33, *p* = .266, η_p_^2^ = 0.077, neither by perceptual load, *F*(1,16) = 0.01, *p* = .943, η_p_^2^ < 0.001, nor an interaction between the two factors, *F*(1,16) = 0.16, *p* = 0.691, η_p_^2^ = 0.010. Time-frequency plots, illustrating contralateral, ipsilateral, as well as contralateral minus ipsilateral power for a broader frequency range (4 – 30 Hz) are available in the supplementary material. The figures show that lateralized activity is restricted to the alpha frequency band. Further, the supplementary material includes a post-hoc analysis, including the factor target eccentricity (inner vs. outer targets, cf. section S2), demonstrating that alpha lateralization was not sensitive to target eccentricity.

**Figure 3.**
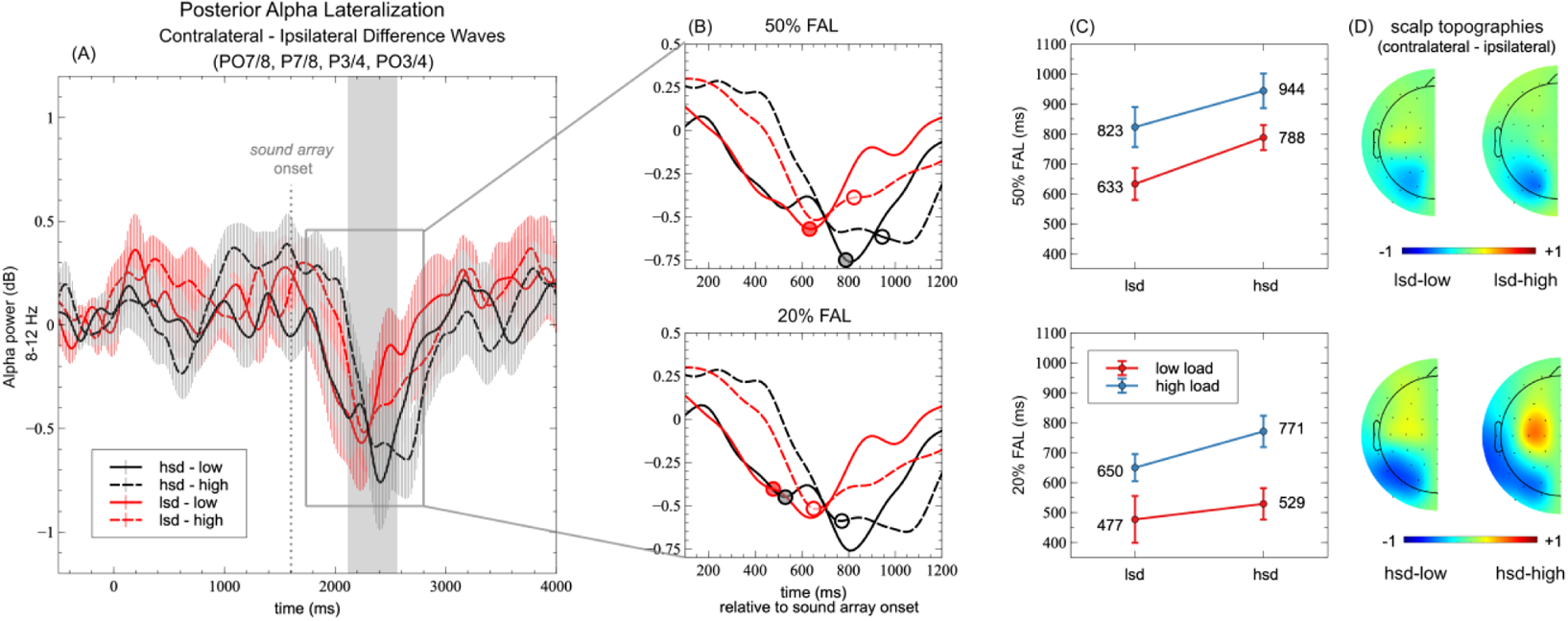
Alpha Power Lateralization. (A) Time course of contralateral minus ipsilateral differences in alpha power across a cluster of parieto-occipital scalp electrodes. The narrow grey rectangle highlights the time window used for statistical analysis of mean alpha lateralization magnitude. Error bars indicate the standard error of the mean. lsd-low = low spatial demand / low perceptual load, lsd-high = low spatial demand / high perceptual load, hsd-low = high spatial demand / low perceptual load, hsd-high = high spatial demand / high perceptual load. (B) A close-up view of the contralateral minus ipsilateral difference waveforms in-between 1700 and 2800 ms (i.e., 100 – 1200 ms relative to sound array onset). The x-axis denotes time (ms) relative to sound array onset. Circles mark the 50% (top) and 20% (bottom) fractional area latency (FAL) measures for each condition. (C) A line plot of the respective 50%-FAL (top) and 20%-FAL (bottom) values, depending on spatial demand and perceptual load. Y-axis values denote FAL relative to sound array onset. Error bars depict the standard error according to Miller et al., 1998 (formula 2) (D) Scalp topographies based on the contralateral minus ipsilateral differences in alpha power in-between 546 to 1061 ms following sound array onset (i.e., the time window used for statistical analyses).

Moreover, to assess the potential impact of horizontal eye movements on alpha lateralization magnitude, the supplementary material includes an additional analysis, including the average lateralized hEOG voltages as a covariate (cf. section S4).

### 3.3 Alpha lateralization onset latencies

To investigate whether the time-course of alpha lateralization was affected by the task demands, we assessed alpha lateralization onset latencies. Figure 3B and C illustrate the points in time where the area under the condition-specific difference curves reaches 20% and 50%, respectively (i.e., the 20% FAL and the 50% FAL). The analysis of fractional area latency (FAL) measures revealed a significant main effect of perceptual load for the 20%-FAL, F_corr_(1,16) = 23.91, *p* < .001, η_p_^2^ = 0.599, and the 50%-FAL, F_corr_(1,16) = 25.39, *p* < .001, η_p_^2^ = 0.613.That is, alpha lateralization emerged earlier in low perceptual load compared to high perceptual load trials (D_20%_ = 208 ms, SE_D-20%_ = 42.21, D_50%_ = 173 ms, SE_D-50%_ = 35.42). Further, a significant main effect of spatial demand was evident for the 20%-FAL, F_corr_(1,16) = 4.84, *p* = .043, η_p_^2^ = 0.232, and the 50%-FAL, F_corr_(1,16) = 9.39, *p* = .007, η_p_^2^ = 0.369, indicating earlier alpha lateralization onset latencies in low spatial demand blocks compared to high spatial demand blocks (D_20%_ = 87 ms, SE_D-20%_ = 39.30, D_50%_ = 138 ms, SE_D-50%_ = 45.25). There were no significant interactions (all F_corr_ < 0.21).

### 3.4 Non-lateralized, posterior alpha power desynchronization

Figure 4 depicts the time-course of posterior alpha power at electrode Pz, separately for each of the four conditions. The analysis revealed a significant main effect of spatial demand, *F*(1,16) = 7.66, *p* = .014, η_p_^2^ = 0.324, reflecting greater alpha ERD (i.e., more negative power) in the high spatial demand condition (M = −2.76 dB, SD = 1.83) compared to the low spatial demand condition (M = −2.14 dB, SD = 1.57). Neither the main effect perceptual load, *F*(1,16) = 3.58, *p* = .077, η_p_^2^ = .18, nor the interaction between spatial demand and perceptual load, *F*(1,16) = 0.64, *p* = .437, η_p_^2^ = 0.04, were significant.

**Figure 4.**
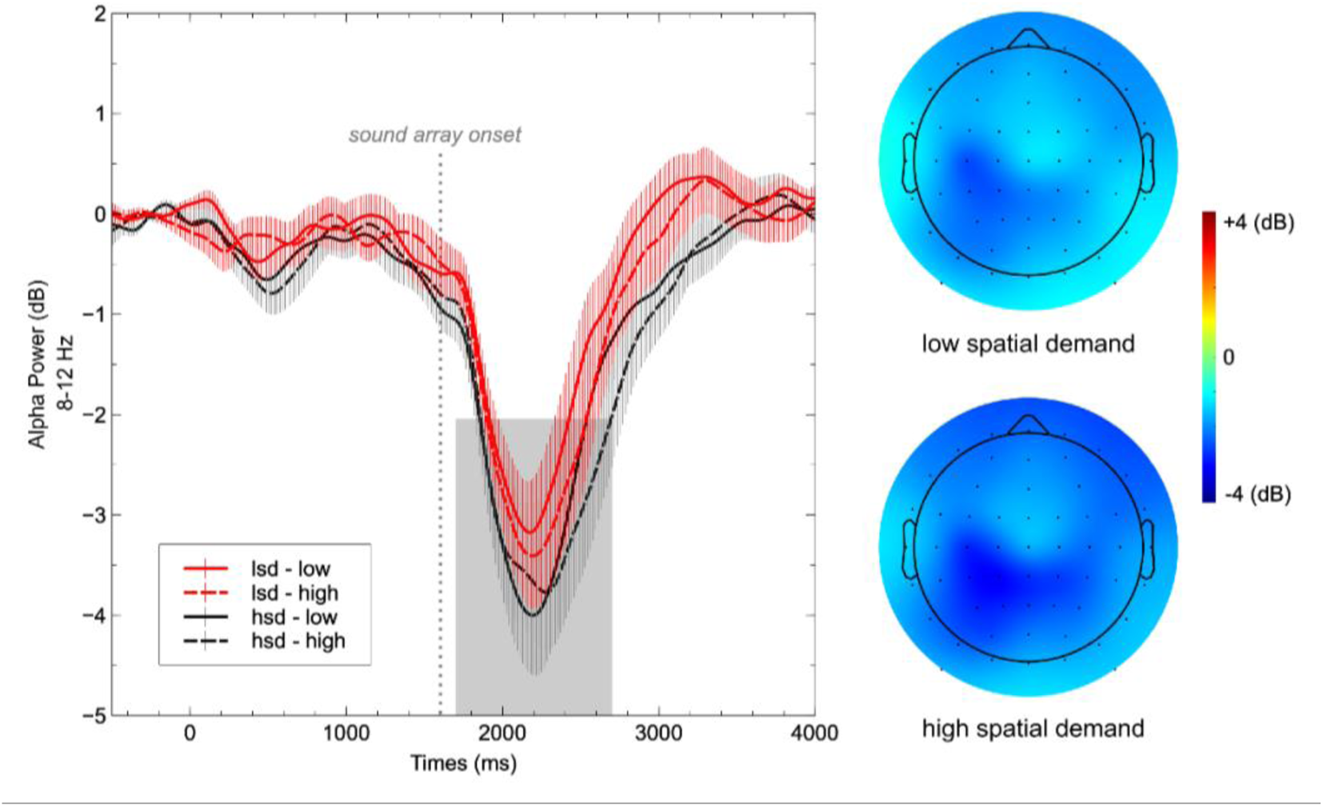
Event-related desynchronization (ERD) of alpha power at Pz. The line plot illustrates the condition-specific averages depending on spatial demand and perceptual load. Error bars indicate the standard error of the mean. lsd-low = low spatial demand / low perceptual load, lsd-high = low spatial demand / high perceptual load, hsd-low = high spatial demand / low perceptual load, hsd-high = high spatial demand / high perceptual load. The grey rectangle indicates the approximate time window used for statistical analysis (i.e., 1698 - 2698 ms relative to target cue onset or 98 - 1098 ms relative to sound array onset). Scalp topographies are based on the average alpha power in the respective analysis time window.

### 3.5 Decoding analysis

We decoded the exact spatial location (i.e., outer-left, inner-left, inner-right, outer-right) of the target sound based on the scalp distribution of alpha-band EEG power. Figure 5 shows the grand average scalp topography for each target location, separately for the two spatial demand conditions (and averaged across the two perceptual load conditions). Figure 6A shows the time-course of decoding accuracy for the low vs. high spatial demand condition, as well as the difference in decoding accuracy between conditions. Decoding accuracy starts to rise above chance level (i.e., 25%) at around 1940 ms in the low spatial demand condition (i.e., 340 ms following sound array onset) and shortly thereafter (around 2100 ms) in the high spatial demand condition. At first, it increases continuously in both spatial demand conditions. In the low spatial demand condition, decoding accuracy reaches a peak at around 2280 ms (i.e., 680 ms post-sound onset) and then gradually decreases throughout the remainder of the response interval; in the high spatial demand condition, decoding accuracy continues to rise beyond the peak in the low spatial demand condition until around ~2440 ms (i.e., 840 ms post-sound onset). While gradually decreasing thereafter, the decoding accuracy remains on a higher level compared to the low spatial demand condition. Toward the end of the response interval (i.e., around 3800 ms), decoding accuracy approaches chance level in both conditions. The cluster mass test revealed that decoding was significantly greater than chance in both spatial demand conditions. We identified significant clusters following sound array onset in each of the two conditions (see Figure 6A, solid green and yellow lines). In the high spatial demand condition, the cluster extends from around 2120 ms to ~3380 ms relative to cue onset (i.e., ~520 – 1780 ms relative to sound array onset, *p* < 10^−4^); in the low spatial demand condition, a first cluster spans a partially overlapping time period in-between ~1940 ms and 2820 ms relative to cue onset (i.e., ~340 – 1220 ms relative to sound array onset, *p* < 10^−4^). In addition, a second, rather late cluster spans the time period from ~3520 – 3740 ms relative to cue onset (i.e., ~1920 – 2140 ms relative to sound array onset, *p* = .036). Note, however, that cluster-based permutation test results should not be used to derive conclusions about the specific onset or offset of a certain effect (Sassenhagen & Draschkow, 2019).

**Figure 5.**
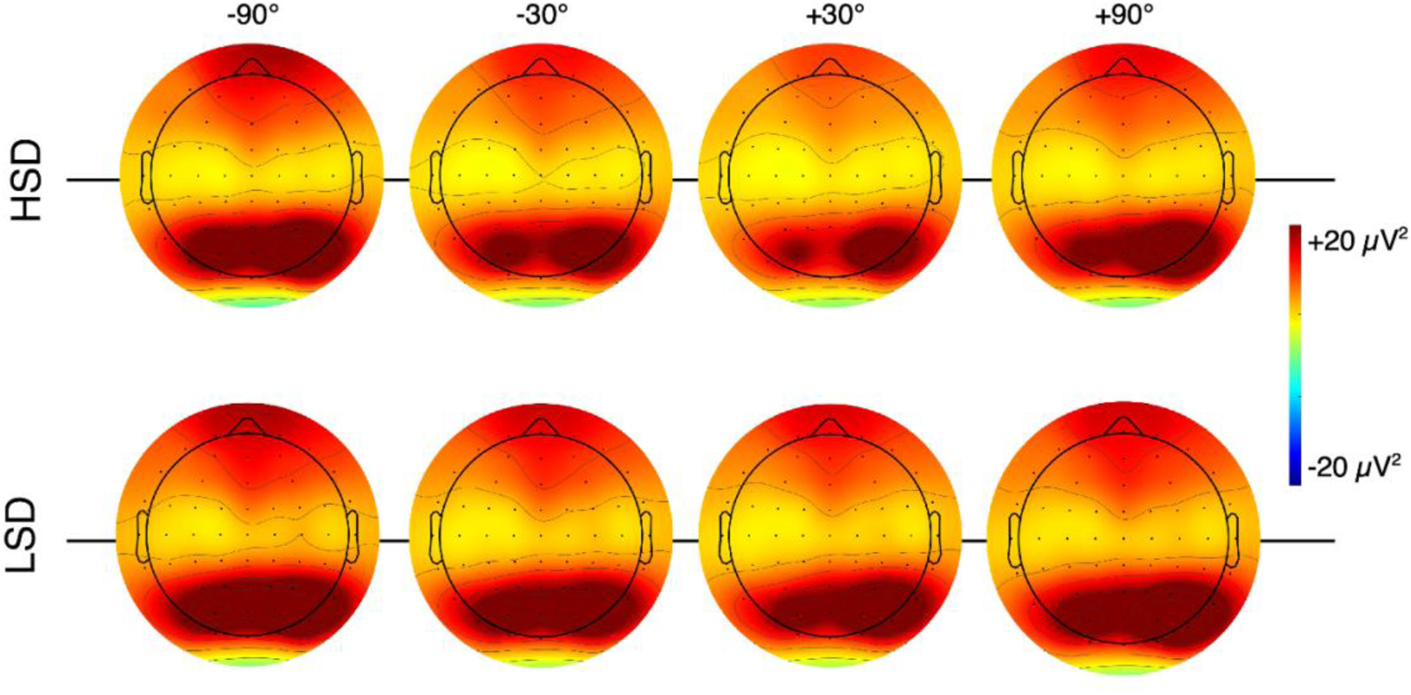
Scalp topographies of instantaneous alpha power for each of the target locations. Alpha power was averaged across a broad time interval following sound array onset (i.e., 320 – 1780 ms post sound array onset), averaged across subjects as well as across the two perceptual load conditions. The top row depicts the scalp topographies for the high spatial demand (HSD) condition, the bottom row shows the low spatial demand (LSD) condition.

**Figure 6.**
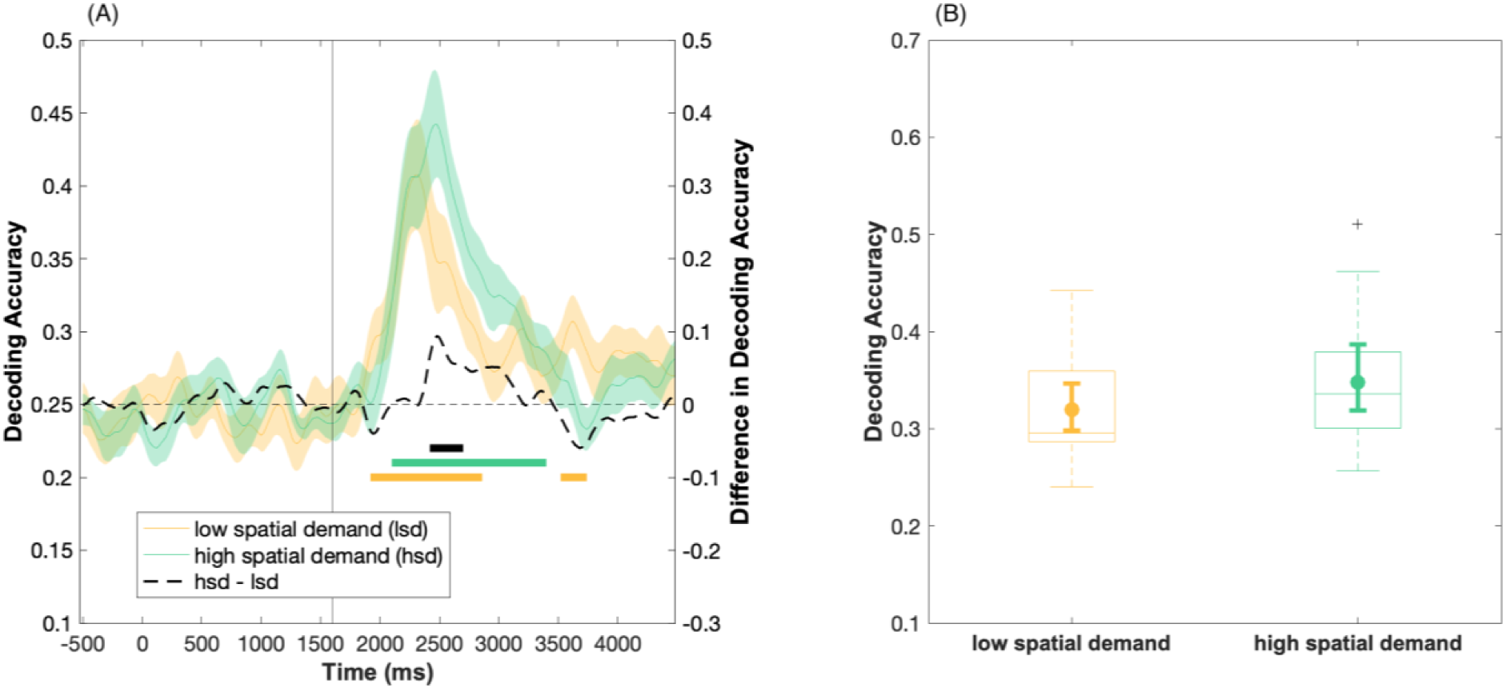
Location decoding based on the multivariate scalp distribution of alpha power. (A) Time-course of the average decoding accuracy results in the low (yellow) and high (green) spatial demand condition, respectively. The colored shading indicates ±1 SEM. Chance-level performance (i.e., 25%) is indicated by the grey dashed horizontal line. The yellow and green solid bars indicate significant decoding of the target location in the low and high spatial demand condition, respectively. The black solid bar denotes significant differences in decoding ability between the low and the high spatial demand condition. Note that only time-points in-between 1600 – 3800 ms were considered in the statistical analysis. The vertical line at x =1600 ms indicates sound array onset. (B) Boxplots refer to the average decoding accuracy in-between 1920 – 3380 ms relative to cue-onset (i.e., 320 – 1780 ms following sound array onset). As per convention, boxplots illustrate the interquartile range and the median. Whiskers extent to the 1.5 times the interquartile range. The superimposed circles show the average decoding accuracy, while the corresponding error bars denote the 95% bootstrap confidence interval of the mean (number of bootstrap samples = 10000).

The black, dashed line in figure 6A illustrates the difference in decoding accuracy between the two spatial demand conditions. A cluster-corrected sign-permutation test indicated significant differences in decoding ability (*p* = .013, one-sided test, cluster extending from ~2420 – 2680 ms relative to cue onset, i.e., ~820 – 1080 ms relative to sound array onset), with higher decoding accuracy in the high spatial demand condition compared to the low spatial demand condition.

Finally, we assessed the overall difference in decoding ability within the post-stimulus period (specifically, within the approximate time-window that resulted in above-chance decoding accuracy within both spatial demand conditions). A one-sided permutation test of the average decoding accuracy between 1920 – 3380 ms (i.e., 320 – 1780 ms relative to sound array onset) consistently revealed a significant difference in decoding accuracy between the spatial demand conditions (*p* = .008, Fig. 6B). Notably, an additional, exploratory decoding analysis based on alpha power at parieto-occipital scalp sites (rather than the whole scalp), returned very comparable results (cf. supplementary material, S3). A supplementary covariate analysis further ruled out that the differences in decoding accuracy can be accounted for by horizontal eye movements (cf. supplementary section S4).

### 3.6 Confusion matrices

To provide more detailed insights into the decoding results, here we show the confusion matrices for each combination of target location and classification response, separately for the high and low spatial demand condition. Figure 7 illustrates the probability of each classification response (i.e., predicted location) for a given stimulus category (i.e., true location), averaged over a broad time window following sound array onset and over participants. In both conditions, the highest probability of classification response is evident at the true location. Interestingly, while neighboring positions receive the most classification errors, the least confusion occurs predominantly between a true location and the position that is in the opposite hemifield and of opposite eccentricity (e.g., left-out vs. right-in).

**Figure 7.**
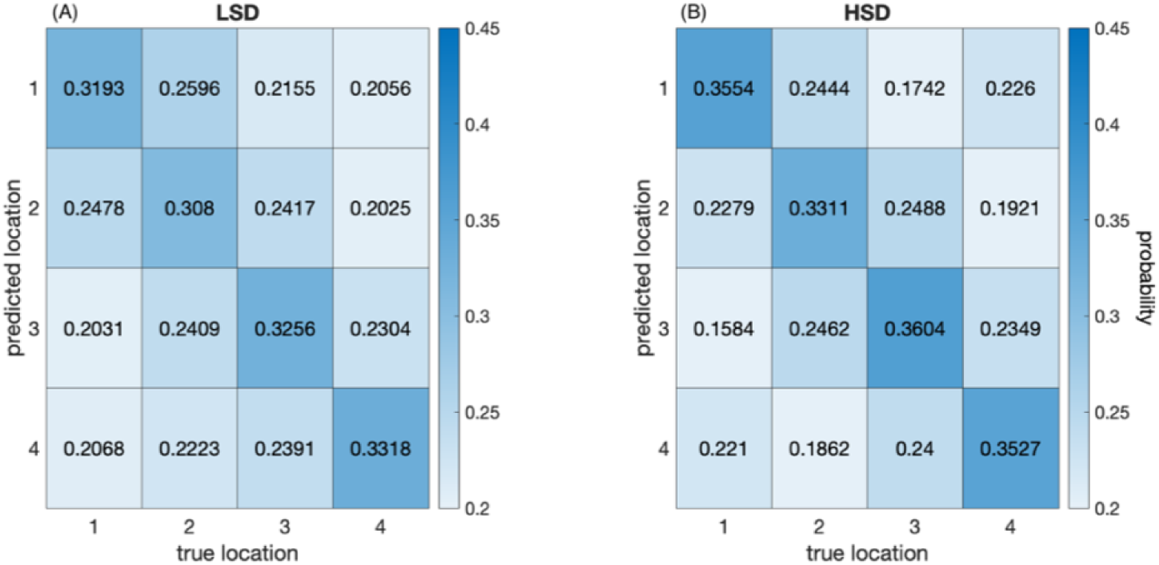
Confusion matrices for the low (LSD, panel A) and high (HSD, panel B) spatial demand condition. Each cell shows the probability of a given classification response (y-axis) for each stimulus position (x-axis), averaged across subjects and across a broad time window following sound array onset (i.e., 1940-3360 ms relative to cue onset or respectively, 340-1760 ms relative to sound array onset). Location labels (1-4) correspond to stimulus locations in their order of occurrence from left to right (i.e., left-out, left-in, right-in, right-out).

### 3.7 Response-locked analyses

As outlined above, we observe both a shift in response latency as well as a shift in alpha-band ERSPs latency across the different conditions. Moreover, the timing of the alpha-band ERSPS seems to coincide well with the timing of the manual responses. This raises the question, to what extent the observed effects could be accounted for by motor response preparation processes (as opposed to perceptual processing). On a similar note, it stands out that the differences in decoding accuracy between the two spatial demand conditions arise relatively late (> 700 ms), which raises the question whether the differences in classification accuracy could be influenced by differences in the response times between the low versus high spatial demand condition. The observations prompted us to perform additional analyses of the response-locked data.

Accordingly, figure 8 illustrates the time-course of alpha-band power, time-locked to the response. It is evident that there is still a pronounced lateralization of alpha power that peaks around the response. However, the analysis revealed neither a modulation by spatial demand or perceptual load, nor an interaction of the two factors (all *F* < 1.80). The analysis of fractional area onset latencies revealed no significant effects, neither for the 20%-FAL (all Fcorr < 1.49), nor for the 50%-FAL (all *F*_corr_ < 0.84).

**Figure 8.**
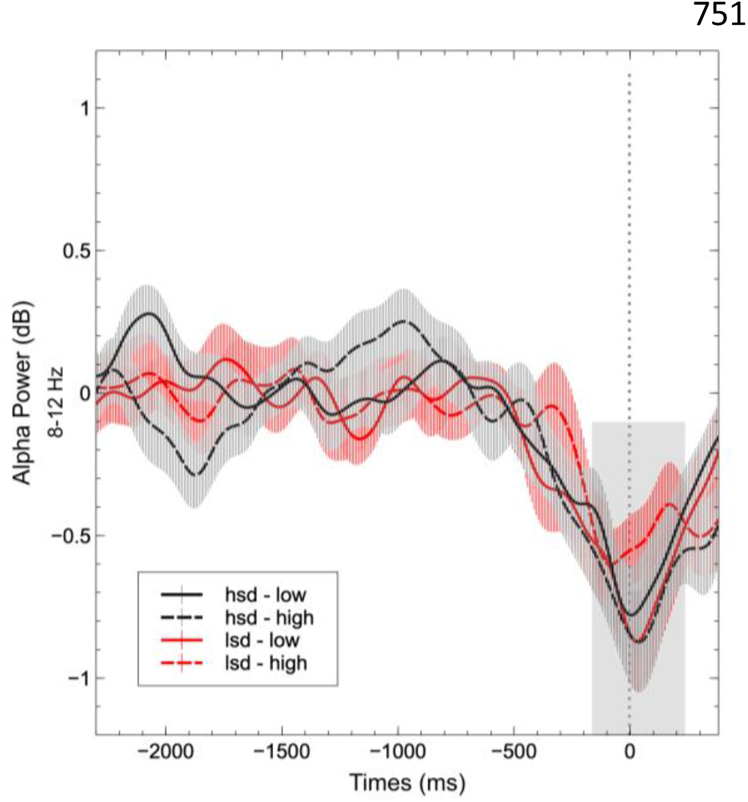
Response-locked alpha power lateralization. The line plot illustrates the condition-specific contralateral minus ipsilateral difference waveforms across a cluster of parieto-occipital scalp electrodes. Error bars indicate the standard error of the mean. The shaded rectangle marks the time window that was used to compute mean alpha power for statistical analyses. The grey dashed line corresponds to the time of the response at x = 0. lsd-low = low spatial demand / low perceptual load, lsd-high = low spatial demand / high perceptual load, hsd-low = high spatial demand / low perceptual load, hsd-high = high spatial demand / high perceptual load.

To investigate whether the differences in decoding accuracy could have been influenced by response time differences, we performed an additional decoding analysis, using the response-locked alpha-band ERSPs as classifier input. Figure 9A illustrate the time course of average decoding accuracy in the low and high spatial demand condition, respectively. In both conditions, decoding accuracy starts to ramp up around −500 ms and then peaks around the response. A cluster-based permutation analysis revealed a significant cluster (*p* < 10^−4^) indicating above-chance decoding accuracy, in both spatial demand conditions (see Figure 9A). In the low spatial demand condition, the cluster ranged from −220 to 360 ms, whereas in the high spatial demand condition, the cluster spans the time period from −340 to 360 ms relative to the response. As mentioned above, cluster-based permutation test results should not be used to derive conclusions about the specific onset or offset of a certain effect (Sassenhagen & Draschkow, 2019).

**Figure 9.**
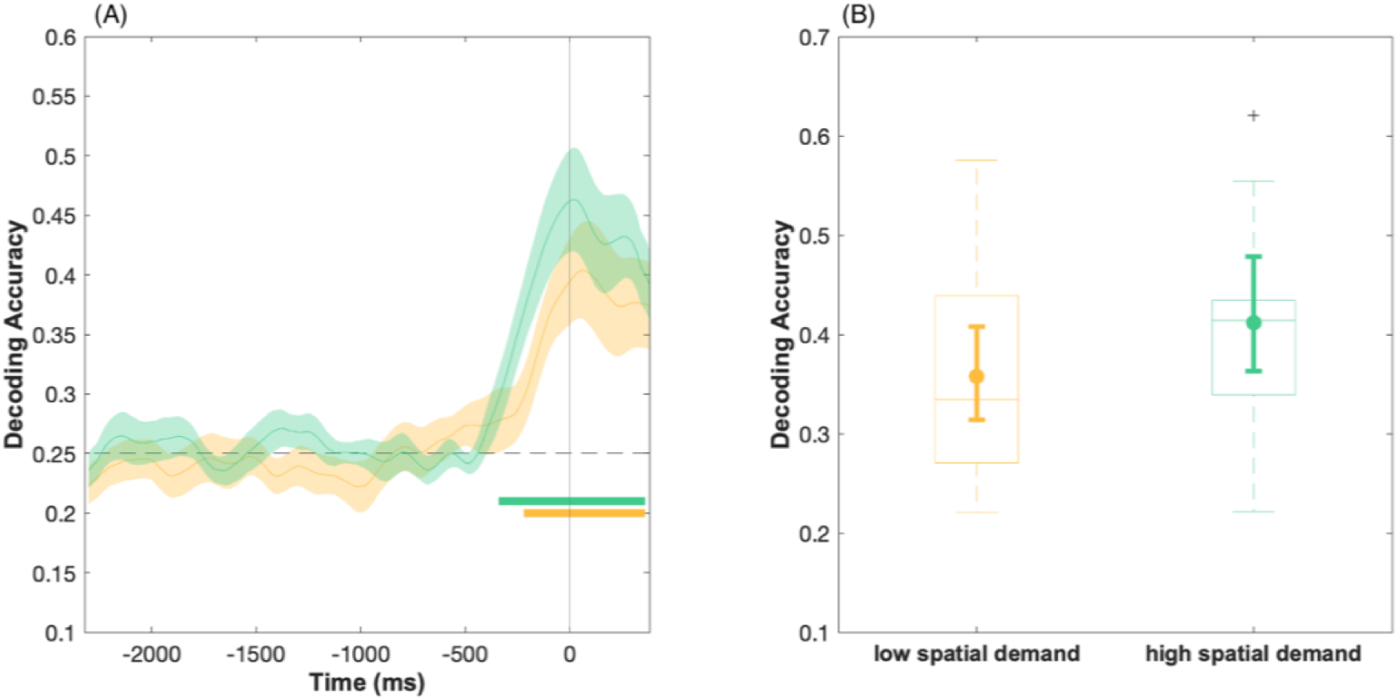
Location decoding based on the multivariate scalp distribution of response-locked alpha-band ERSPs. (A) Time-course of the average decoding accuracy in the low (yellow) and high (green) spatial demand condition, respectively. The colored shading indicates ±1 SEM. Chance-level performance (i.e., 25%) is indicated by the grey dashed horizontal line. The yellow and green solid bars indicate significant decoding of the target location in the low and high spatial demand condition, respectively. Note that only time-points in-between −500 and 380 ms were considered in the statistical analysis (360ms corresponds to the last post-response sampling point in the ERSP waveforms). The vertical line at x =0 ms indicates the response. (B) Boxplots refer to the average decoding accuracy in-between −340 – 360 ms relative to the response. As per convention, boxplots illustrate the interquartile range and the median. Whiskers extent to the 1.5 times the interquartile range. The superimposed circles show the average decoding accuracy, while the corresponding error bars denote the 95% bootstrap confidence interval of the mean (number of bootstrap samples = 10000).

Finally, we computed the average decoding accuracy in-between a broad time window that resulted in significant within-condition decoding across both conditions (i.e., −340 to 360 ms). A one-sided permutation test reveals that on average classifier performance in the high spatial demand condition still significantly exceed classification accuracy in the low spatial demand condition (*p* = .005).

## 4 Discussion

Sensory stimuli and behavioral demands are constantly subject to change, requiring the attentive brain to adapt its response to accommodate to those changes. In this study, we investigated the effects of varying perceptual load and spatial demand in a sound localization task on post-stimulus alpha-band oscillations. The notion that alpha-band oscillations track the currently attended location in a spatially fine-tuned manner is relatively undisputed. However, what remains more elusive is to what degree this spatial specificity depends on the current task demands. Here, we demonstrate that the amount of spatial information reflected in the multivariate scalp distribution of alpha power increases when the task requires a precise sound localization (i.e., indicating the exact stimulus location) compared to when a rather coarse localization judgment is required (i.e., indicating the hemifield). In contrast, these task demand-dependent modulations were not captured by the magnitude of univariate parieto-occipital alpha lateralization. Rather, the time course of alpha power lateralization varied with the task demands.

Behaviorally, the pattern of results was consistent with the well-established observation that the detection of a target sound in a cocktail-party scenario suffers from additional concurrent stimuli in the auditory scene (Brungart & Simpson, 2007; Brungart, Simpson, Ericson, & Scott, 2001; Ericson, Brungart, & Simpson, 2004; Klatt et al., 2018b). Accordingly, in the present study, participants’ responses were slower and less accurate when the sound array contained four (high perceptual load) instead of just two sounds (low perceptual load). In terms of sound localization accuracy, this difference was even more pronounced when they were asked to report the exact target location (high spatial demand) rather than the target hemifield (low spatial demand). Certainly, the present set size effect cannot be completely disentangled from the effects of energetic masking due to the acoustic overlap between the competing sound sources (cf. Murphy, Spence, & Dalton, 2017). However, most critical for the intended EEG analysis was the manipulation of spatial demand. As expected, indicating the exact sound location was more challenging (i.e., slower and less accurate) than simply determining whether the target was present in the left or right hemispace. Nevertheless, subjects still managed to perform clearly above chance level (i.e., on average > 80% correct).

### 4.1 Decoding of auditory covert attention based on alpha power modulations

The main question of the present study was: Is the difference in spatial task demands also reflected in the neural signal? Strikingly, while the classifier could reliably decode the precise target location in both spatial demand conditions, the amount of spatial information reflected in the scalp distribution of alpha-band power was higher under high spatial demand. It should be emphasized that in both spatial demand conditions, participants were presented with the exact same trials (although in randomly shuffled order). This rules out that differences between conditions were caused by bottom-up perceptual factors. Further, the confusion matrices show most classification errors for neighboring positions and the least confusion between the true target location and the location that is both within the other hemifield and on the opposite side (e.g., left-out vs. right-in). This supports the assumption that in auditory scene analysis the relative location between sounds is coded on a neural level, rather than the mere stimulus position (cf. Shiell et al., 2018).

The present results extend our previous work, using an analogous auditory search task design, where we demonstrated that the presence of auditory post-stimulus alpha lateralization was dependent on the task-relevance of spatial information. Specifically, Klatt et al. (2018b) showed that alpha lateralization was absent in a simple sound detection task (i.e., when spatial location was completely irrelevant to the task), whereas it reliably indicated the attended location when participants were asked to localize the target. Here, we show that post-stimulus alpha oscillations are not only sensitive to such coarse manipulations of spatial relevance, but rather – when considering the multivariate activity patterns – also capture fine-grained adaptions to the required degree of spatial specificity. However, the curves reflecting decoding accuracy in the low and high spatial demand conditions do not diverge until about 660 ms following sound array onset; in addition, statistically significant differences in decoding accuracy were limited to a relatively late time-window (i.e., > 800 ms following sound array onset; cf. Figure 5A). In contrast, general decodability of spatial location increases above chance level shortly after sound array onset and persists well into the response interval. This suggests that, even though the spatial demand conditions were blocked (i.e., participants knew beforehand which spatial specificity would be required), it took several hundred milliseconds to evoke changes in spatial specificity of the underlying alpha power signal. Such long latencies have also been reported with respect to voluntary adaptions of the alpha-power signal in a visual spatial cueing study paradigm, requiring participants to adopt either a narrow or a broad focus of attention in anticipation of an upcoming search array (Feldmann-Wüstefeld & Awh, 2019). In the latter study, Feldmann-Wüstefeld and Awh (2019) computed spatially selective channel tuning functions (CTF) based on the topography distribution of alpha power and assessed their slope as a measure of spatial selectivity. Notably, differences in the CTF slopes between the narrow-focus cue and the broad-focus cue only emerged at timepoints > 500 ms following cue onset. Considering the response times in the present study, the question arises, whether the observed differences in decoding accuracy could be influenced by response time differences between the two spatial demand conditions. However, a decoding analysis based on the response-locked alpha-band ERSPs still revealed, on average, greater decoding accuracy in the high compared to the low spatial demand condition. This clearly shows that the condition differences in the stimulus-locked analysis do not solely rely on differences in response time.

Critically, the present results add to these previous findings in several ways: First, we demonstrate that just like preparatory attention is finely tuned and spatially sharpened depending on the task demands (Feldmann-Wüstefeld & Awh, 2019; Voytek et al., 2017), the ongoing attentional processing following search array onset is dynamically modulated depending on the required spatial specificity of the task. Further, the present findings complement a growing body of evidence, supporting the assumption that modulations of alpha oscillations represent a ubiquitous top-down controlled mechanism of spatial attention that plays a role across different attentional domains as well as across sensory modalities. Notably, the pattern that decoding accuracy increases if a more precise spatial judgment is required did fully reproduce when using only parieto-occipital channels as classifier input (cf. supplementary material). This suggests that most information that contributes to classification performance, and critically, to the difference in decoding accuracy between conditions, is present at posterior electrode sites. Overall, this is in line with the notion that parieto-occipital cortex subserves a supramodal neural circuit for spatial attention (Popov, Gips, Weisz, & Jensen, 2021). Although, to be fair, the present analysis does not provide precise insights into the features that drive decoding performance in the two conditions. It remains possible that there are more subtle differences in the features (e.g., the mixture of electrodes) that contribute to the classification accuracy in the two spatial demand conditions.

In principle, the notion that attention can improve the information content of a neural code is not novel. In fact, it is well-established that attending to a spatial position or a relevant feature increases single-neuron firing rates in primary and extrastriate visual areas and can result in changes in the size and position of spatial receptive fields (reviewed by (Sprague, Saproo, & Serences, 2015). In the auditory domain, physiological recordings in cats (Lee & Middlebrooks, 2011) revealed similar sharpening of spatial tuning in auditory cortex (i.e., A1) when the animal engaged in a spatial task compared to an off-task “Idle” condition and a non-spatial periodicity detection task (for similar findings in human A1 see van der Heijden, Rauschecker, Formisano, Valente, & de Gelder, 2018). Hence, along with previous studies in the visual modality (Feldmann-Wüstefeld & Awh, 2019; Voytek et al., 2017), the present results extend these findings, showing that such “sharpening” of neural activity occurs not only in tuning functions of single neurons, but is also evident in the adaption of population-level activity patterns.

### 4.2 Alpha power lateralization as a temporally resolved signature of target processing

In addition to the multivariate decoding analysis, we also analyzed alpha lateralization following sound array onset as a ‘classical’ univariate measure of attentional orienting (e.g., Ikkai, Dandekar, & Curtis, 2016). In the present study, alpha lateralization magnitude did neither vary with perceptual load or spatial demand. The former observation replicates results of a previous study (Klatt et al., 2018b), finding no evidence for differences in alpha lateralization magnitude between a low-load (i.e., two-sound array) and a high-load (i.e., four-sound array) auditory search condition. In contrast, Bacigalupo and Luck (2019) reported that target-elicited alpha lateralization in a visual search paradigm tended to increase with greater task difficulty. Thus, the authors speculate that alpha lateralization might reflect effort rather than target selection. The present findings do not seem to bolster this claim: Both the behavioral data as well as the complementary analysis of non-lateralized posterior alpha power indicate that task difficulty and required cognitive resources increased with greater spatial demand. Yet, alpha lateralization magnitude was unaffected by the experimental manipulation. An additional study by Wang, Megla, and Woodman (2021) corroborates the present results, showing that the magnitude of stimulus-induced alpha lateralization remains unaffected by an increase in the difficulty of attentional selection (e.g., through higher distractor numerosity), while global, non-lateralized posterior alpha power suppression did increase with distractor set size (experiment 1 and 2) and with greater distance between the items (experiment 3). Although, in addition to a general caveat about the interpretation of non-significant effects, it should be noted that with the present sample size, we were likely limited to detect medium-sized effects (see also section 2.2).

Nonetheless, the present findings do substantiate the notion that post-stimulus (or target-elicited) alpha lateralization presents an active signature of target processing in both visual (Bacigalupo & Luck, 2019) as well as auditory search (Klatt et al., 2018b). Bacigalupo and Luck (2019) further disscociate alpha lateralization from a well known ERP-signature of target individuation (i.e., the N2pc), suggesting that alpha lateralization reflects a long-lasting and ongoing attentional processing of the target. Although we do not investigate ERP correlates in the present study, a closer look at the time-course of alpha lateralization supports this assumption: on average, alpha lateralization persist beyond and in fact peaks around the time participants make their response. Different temporal characteristics of N2ac (an auditory analogue of the visual N2pc commponent Gamble & Luck, 2011) and alpha lateralization have recently also been observed in response to shifts of auditory attention between relevant talkers in a simulated cocktailparty scenario (Getzmann, Klatt, Schneider, Begau, & Wascher, 2020), corroborating the notion that the EEG measures reflect different attentional processes (see also Klatt et al., 2018b).

Contrary to alpha lateralization magnitude, alpha lateralization onset latency was linked to task demands. Specifically, alpha laterization emerged around 170 ms earlier (50%-FAL) in the less demanding low perceptual load condition relative to the high perceptual load condition and ~140 ms earlier (50%-FAL) in the low spatial demand condition relative to the high spatial demand condition. Overall, the observed modulations of alpha lateralization onset latency are in line with a previous visual search study (Foster et al., 2017), showing that the onset of alpha-based CTFs varied with reaction times as well as search difficulty.

That the latency differences reported by Foster et al. (2017) were much larger (i.e., differences of up to 440 ms) could be attributed to the fact that their search conditions differed more strongly (e.g., distractors were all identical vs. heterogenous). In sum, the present findings corroborate the claim that attentional modulations of alpha power not only track the location of covert spatial attention, but also the time-course (i.e., the latency) of post-stimulus attentional processing.

Moreover, it is striking that alpha lateralization is still evident in the response-locked ERSPs – notably, of similar magnitude as in the stimulus-locked ERSPs – whereas the modulations of ERSP latency were not present anymore. While this rules out that the observed effects can be better accounted for by motor preparation processes than by perceptual processing, together with the response-locked decoding analysis, it does support the notion that alpha oscillations might be closely related to the transfer of spatially-specific information in a response-specific format (Klatt et al., 2018a).

Finally, the apparent difference between univariate and multivariate measures of alpha power highlights the potential of multivariate decoding for the study of neurocognitive mechanisms. Similarly, when performing a univariate analysis of alpha power, Voytek et al. (2017) did not capture the fine-grained differences in the allocation of attention (depending on the spatial certainty of a cue) that were evident in the multivariate topography of alpha power. Taken together, this illustrates the increased sensitivity of multivariate decoding techniques to reveal complex dynamics that are present in the combined signal across the scalp (Hebart & Baker, 2017).

## 5 Conclusion

In conclusion, our results show that the spatial specificity of post-stimulus alpha-band oscillations can be finely adapted depending on the spatial demands of the task. Notably, this task-dependent adaptation was only evident in the multivariate distribution of the alpha-band signal, whereas the magnitude of univariate parieto-occipital alpha lateralization did not capture those variations in perceptual load and spatial demand. Rather, alpha lateralization onset latency varied with the difficulty of the task, suggesting that the time-resolved modulation of post-stimulus alpha lateralization captures differences in the efficiency of post-attentional processing. These findings improve our understanding of the functional role of alpha oscillations for the ongoing attentional processing of complex auditory scenes and provide new insights into the attentional mechanisms underlying top-down adaptions to changing task demands.

## 6 Competing Interests

Declarations of interest: none.

## 7 Author contributions: CRediT authors statement

**Laura-Isabelle Klatt:** Conceptualization, Formal analysis, Investigation, Writing – Original Draft, Visualization, Project administration **Stephan Getzmann:** Conceptualization, Writing – Review & Editing, Supervision **Daniel Schneider:** Conceptualization, Writing – Review & Editing, Supervision

## 8 Acknowledgements

The authors would like to thank Peter Dillmann for the technical implementation of the stimulus presentation software and Tobias Blanke for technical support. Further, the authors are grateful to the lab staff, in particular to Pia Deltenre and Barbara Foschi, and their team of student assistants for their help with data acquisition. In addition, many thanks go to Gi-Yeul Bae and Steven Luck as well as Michael J. Wolff and colleagues for publicly sharing their code (Bae & Luck, 2018; Wolff et al., 2017), which served as the basis for the present decoding and cluster permutation analyses.

## 9 Funding

This research did not receive any specific grant from funding agencies in the public, commercial, or not-for-profit sectors.

## SUPPLEMENTARY MATERIALS

### S1. Time-Frequency Plots

Supplementary figure 1 illustrates contralateral, ipsilateral, as well as contralateral minus ipsilateral power at a cluster of posterior electrodes (PO7/8, P7/8, P3/4, or PO3/4) for a frequency range of 4 to 30 Hz separately for each condition. The figure confirms that lateralization effects are mostly limited to the alpha frequency range (8-12 Hz).

**Figure 1.**
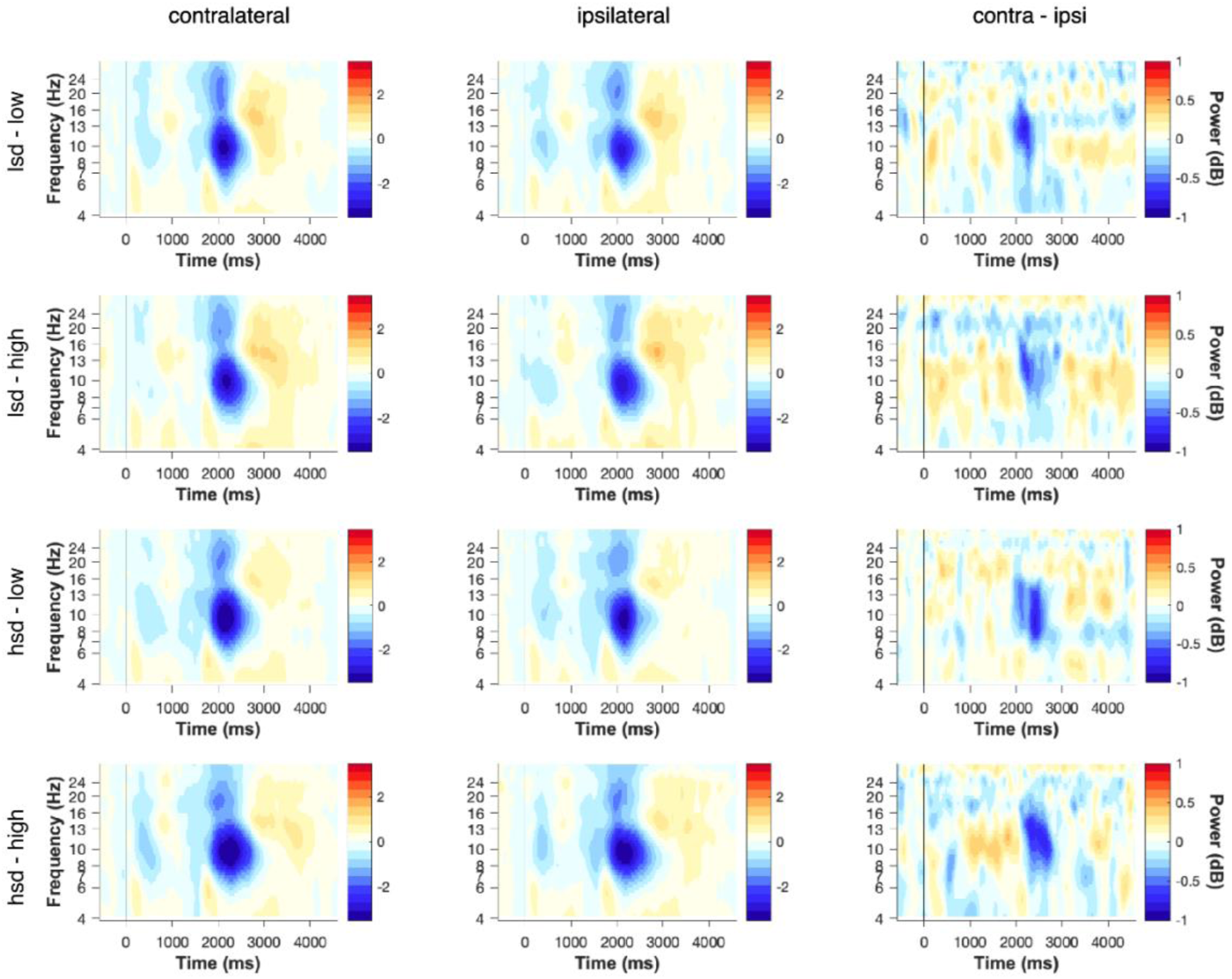
Time-frequency plots. Power is depicted for a frequency range of 4 to 30 Hz at electrodes contralateral (left column) and ipsilateral (middle column) to the target location as well as for the contralateral minus ipsilateral (right column) differences. The conditions are abbreviated as follows: lsd-low = low spatial demand / low perceptual load, lsd-high = low spatial demand / high perceptual load, hsd-low = high spatial demand / low perceptual load, hsd-high = high spatial demand / high perceptual load.

### S2. Alpha lateralization magnitude and target eccentricity

Deng, Choi, and Shinn-Cunningham (2020) have previously reported that alpha lateralization is greater when attention was directed to locations further away from the central position. In the present study, targets could be likewise presented in rather close proximity to central fixation (± 30°) or at a greater distance (± 90°). Thus, in the present study, the question arises whether modulations of alpha lateralization magnitude depending on target eccentricity (inner vs. outer targets) are only present in the high spatial demand condition, while alpha lateralization might differentiate between those positions in the low spatial demand condition (or does so to a lesser extent). Hence, we computed alpha lateralization magnitude separately for inner (± 30°) and outer (± 90°) targets. Then, we performed post-hoc paired-sample *t*-tests to contrast the difference in alpha lateralization magnitude between inner minus outer targets for the LSD versus HSD condition (separately for the low and high perceptual load condition). Neither in the low perceptual load condition *t*(16) = −0.11, *p* = .915, *p*_adj_ = 1.372, nor in the high perceptual load condition, *t*(16) = −1.47, *p* = .160, *p*_adj_ = .480, did we find that the difference in alpha lateralization magnitude between inner vs. outer targets differed between the low and the high spatial demand condition.

### S3. Decoding based on alpha power at parieto-occipital scalp sites

In previous studies that applied alpha-band decoding, results have been shown to be virtually identically for analyses including all vs. only posterior electrodes (e.g., van Moorselaar et al., 2018), suggesting that most (or even all) relevant information that contributes to decoding performance is represented in posterior electrode sites. Hence, we performed an additional, exploratory decoding analysis, using only parieto-occipital scalp sites as input to the classifier. Otherwise, all parameters in the decoding analysis were used, as described in the main analysis. Specifically, alpha power at electrodes P7, P3, P1, Pz, P2, P4, P8, PO7, PO3, POz, PO4, PO8, O1, Oz, O2, PO9, and PO10 was used as input to the classifier. Supplementary figure 3A shows the resulting time-course of decoding accuracy as well as the difference between the low and high spatial demand condition. In addition, panel B illustrates the average decoding accuracy per condition.

**Figure 2.**
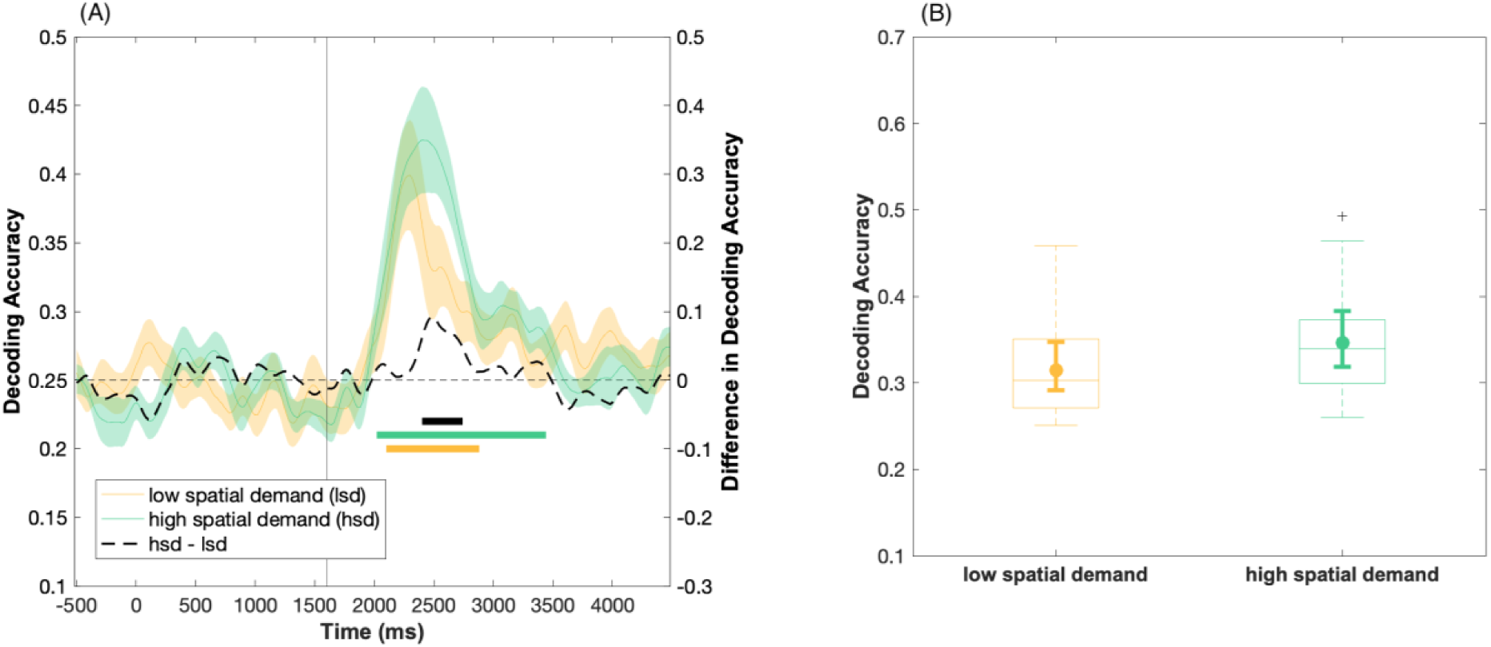
Location decoding based on the scalp distribution of alpha power at posterior electrode sites. (A) Time-course of the average decoding accuracy results in the low (yellow) and high (green) spatial demand condition, respectively. The colored shading indicates ±1 SEM. Chance-level performance (i.e., 25%) is indicated by the grey dashed horizontal line. The yellow and green solid bars indicate significant cluster of decoding accuracy in the low and high spatial demand condition, respectively (cluster-based permutation, all *p* < 10^−4^). The black solid bar denotes significant differences (cluster-corrected sign-permutation test, one-sided, *p* = .006) in decoding ability between the low and the high spatial demand condition. Note that only time-points in-between 1600 – 3800 ms were considered in the statistical analysis. (B) Boxplots refer to the average decoding accuracy in-between 2040 - 3400 ms relative to cue-onset (i.e., 440 – 1800 ms following sound array onset). The latter time window includes the approximate time interval that resulted in significant within-condition decoding for both conditions. As per convention, boxplots illustrate the interquartile range and the median. Whiskers extent to the 1.5 times the interquartile range. The superimposed circles show the average decoding accuracy, while the corresponding error bars denote the 95% bootstrap confidence interval of the mean (number of bootstrap samples = 10000). A one-sided permutation test revealed a significant difference in the overall decoding ability between the low and high spatial demand condition, *p* = .001.

### S4. Control analyses: eye movements

We evaluated the epoched data at horizontal EOG (hEOG) electrodes. To obtain ERPs, the continuous EEG data was segmented into epochs, ranging from −1000 to 4500 ms relative to sound cue onset. For baseline correction, the 200 ms time interval prior to sound cue onset was used (i.e., −200 to 0 ms). Trials classified as premature responses (i.e., with response times < 200 ms) were removed. Otherwise, no preprocessing was performed.

Supplementary figure 3 depicts the contralateral versus ipsilateral voltages at hEOG electrodes LO1/LO2 relative to the target position. The ERP at horizontal EOG electrodes clearly indicates typical lateralized voltage differences, depending on the target position. Thus, despite the instruction to fixate the central LED, on at least a subset of trials, saccades toward the target sounds were present. Figure 3 also shows that saccades were more pronounced in the high spatial demand condition.

**Figure 3.**
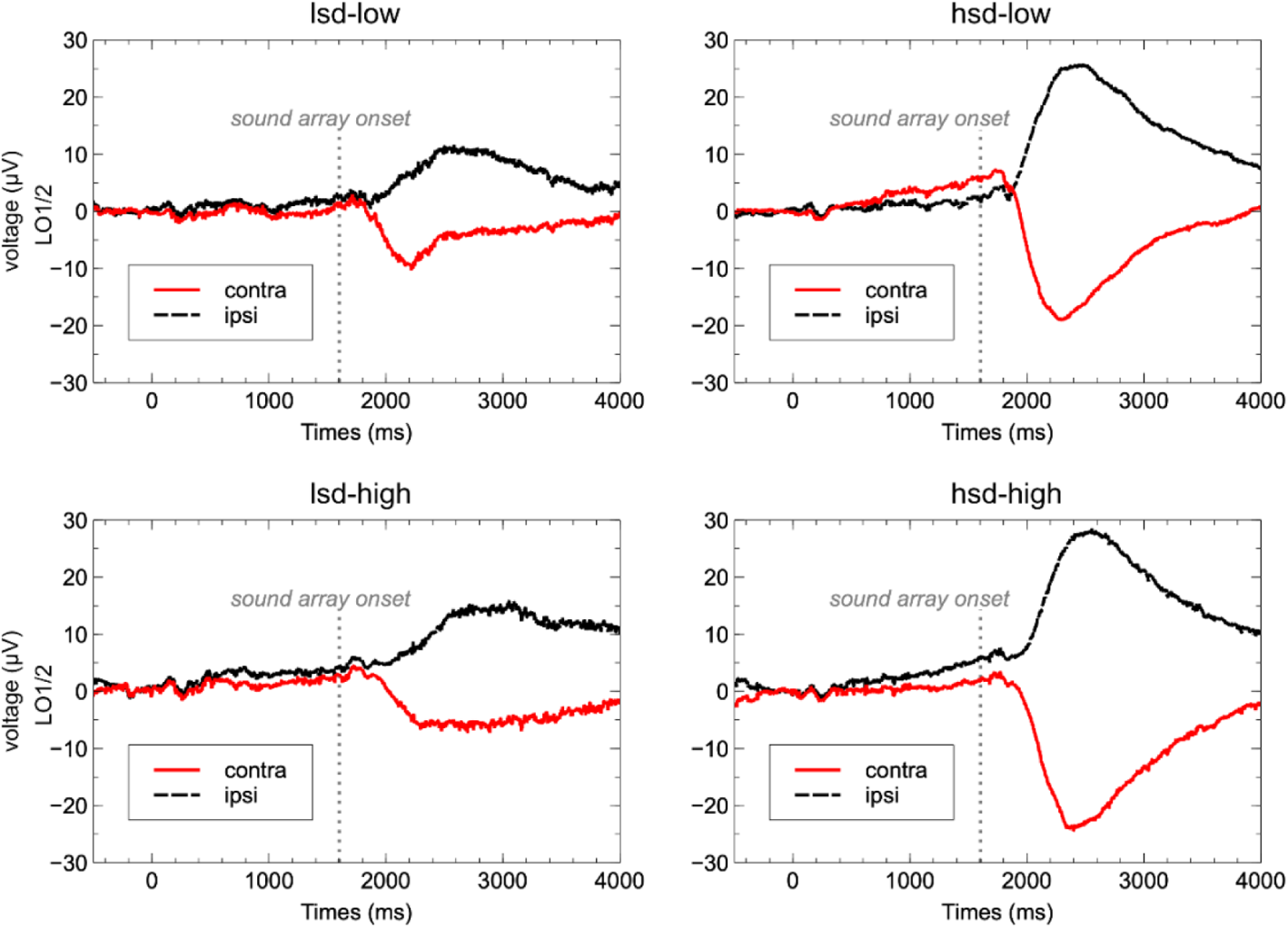
Contralateral versus ipsilateral voltages at lateral EOG electrodes LO1 and LO2 relative to the target position for each of the four conditions.

To assess the potential impact of horizontal eye movements on alpha lateralization magnitude, we conducted an analysis of covariance, including spatial demand (high vs. low), perceptual load (high vs. low), and asymmetry (contra vs. ipsi) as within-subject factors and the average lateralized hEOG as a covariate. Specifically, to obtain the average lateralized hEOG voltages, the contralateral minus ipsilateral waveforms for each subject were averaged across all conditions. In the resulting average waveforms, mean amplitude was measured in-between 2000 – 3000 ms post cue onset (i.e., 400 – 1600 ms post-sound array). Mean alpha power (using the same time window as reported in the main manuscript) served as the dependent variable. We found that the covariate (i.e., saccades) was not significantly related to the magnitude of alpha-band lateralization, as indicated a non-significant interaction of the factor saccades and asymmetry, *F*(1,15) = 0.21, *p* = .656, η_p_^2^ = 0.014. Importantly, the main effect of asymmetry remained significant, *F*(1,15) = 8.69, *p* = .010, η_p_^2^ = 0.367, after controlling for the effect of saccades. This suggests that the overall presence of alpha lateralization is not affected by the occurrence of horizontal eye movements. Further, none of the interactions, involving the factor saccades turned out to be significant, all *p* > .10.

Further, to assess the potential impact of eye movements on our decoding results, we performed an exploratory decoding analysis, using the horizontal (LO1/LO2) and vertical (IO1/IO2) EOG channels as input. The decoding analysis was performed as described in the main manuscript, with the following exception: Given that all trials (except for individual premature responses, target-absent trials and incorrectly answered trials) served as input to the decoding analysis, a five-fold (rather than a three-fold) cross validation was performed. Supplementary figure 4 illustrates the time-course of decoding accuracy for the low and high spatial demand condition, respectively. In both conditions, it was possible to decode target location based on hEOG input (all *p* < 10^−4^). However, in contrast to the main decoding analysis based on the whole-scalp topography of alpha power, we did not find a significant difference in decoding accuracy between the high vs. low spatial demand condition (cluster-corrected sign-permutation test, *p* = .22). In addition, overall, decoding accuracy was lower for hEOG based decoding compared to alpha power decoding, which matches an observation also made by Popov, Gips, Weisz, & Jensen (2021). Thus, even though it appears that saccades are more pronounced in the high spatial demand condition, this does not account for the higher decoding accuracy in the high spatial demand condition that becomes apparent in the alpha-based decoding.

**Figure 4.**
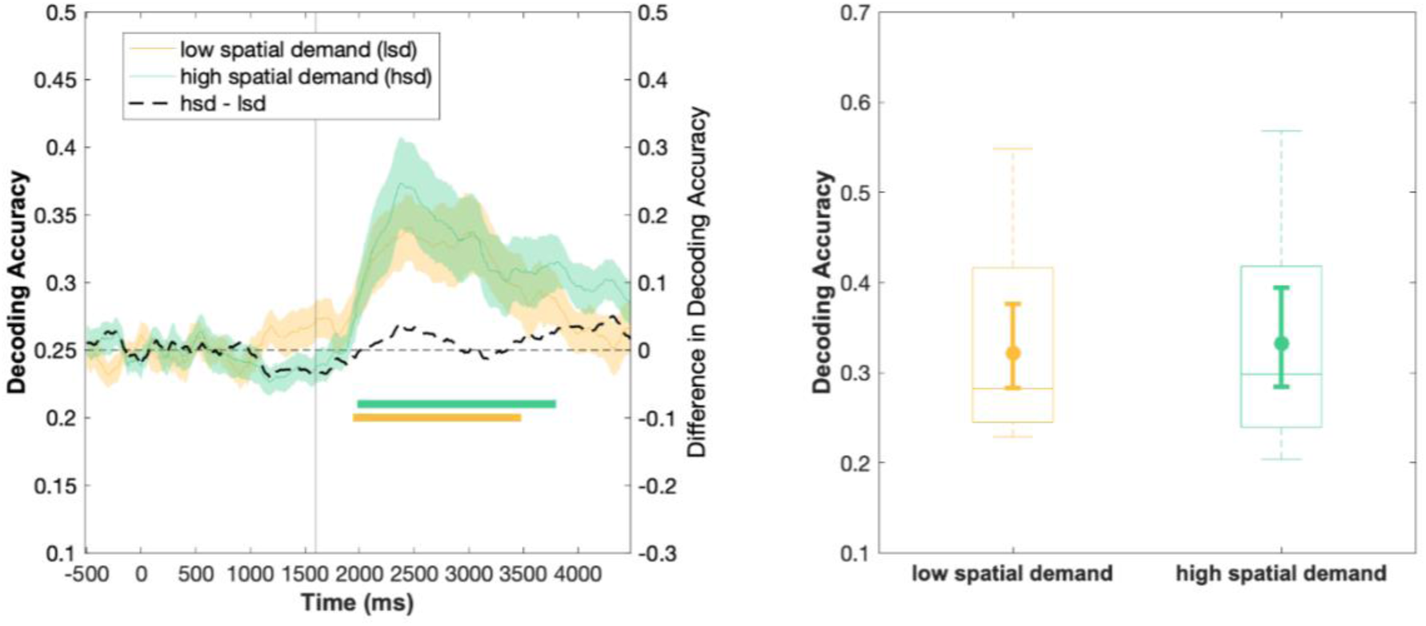
Location decoding based on the ERP signal at horizontal and vertical EOG electrodes. (A) Time-course of the average decoding accuracy results in the low (yellow) and high (green) spatial demand condition, respectively. The colored shading indicates ±1 SEM. Chance-level performance (i.e., 25%) is indicated by the grey dashed horizontal line. The yellow and green solid bars indicate significant decoding of the target location in the low and high spatial demand condition, respectively. The black solid bar denotes significant differences in decoding ability between the low and the high spatial demand condition. Note that only time-points in-between 1600 – 3800 ms were considered in the statistical analysis. (B) Boxplots refer to the average decoding accuracy in-between 1960 – 3500 ms relative to cue-onset (i.e., 300 – 1900 ms following sound array onset). As per convention, boxplots illustrate the interquartile range and the median. Whiskers extent to the 1.5 times the interquartile range. The superimposed circles show the average decoding accuracy, while the corresponding error bars denote the 95% bootstrap confidence interval of the mean (number of bootstrap samples = 10000).

Finally, to follow up more closely on whether the apparent differences in hEOG asymmetry between conditions (and between subjects) systematically covary with the effect of spatial demand on alpha-based decoding accuracy, we performed a post-hoc ANCOVA: The average alpha-band based decoding accuracy in-between 1920 – 3380 ms (i.e., broad time window that resulted in significant within-condition decoding across both spatial demand conditions) served as a dependent variable, spatial demand (low vs. high) served as a within-subject factor, and the difference in hEOG lateralization (2000 – 3000 ms post cue onset) between the spatial demand conditions (hsd – lsd) was included as a covariate (referred to as factor ‘saccades’). The covariate was not significantly related to decoding accuracy, *F*(1,15) = 0.80, *p* = .384, η_p_^2^ = 0.051. Critically, the main effect of spatial demand was still significant after controlling for effect of saccades, *F*(1,15) = 5.33, *p* = .036, η_p_^2^ = 0.262. The interaction between saccades and spatial demand was not significant, *F*(1,15) = 0.01, *p* = 0.922, η_p_^2^ = 0.001.

Although we need to consider that a non-significant association between a covariate (i.e., hEOG asymmetry) and a dependent variable (e.g., decoding accuracy or alpha lateralization magnitude) does not proof that there is no such relationship, what is critical, is the fact that the main effects of asymmetry and spatial demand, respectively, remain significant after controlling for the effects of saccades. Although, it should be noted that a caveat applies since we are analyzing the same data with and without considering a covariate; thus, the presented covariate analyses should be considered exploratory.

Taken together, the above presented control analyses reassure that the most critical finding – namely, the greater decoding accuracy for the high spatial demand condition – is not based on systematic difference in eye movements. Although we cannot be perfectly sure that the within-condition decoding of target location has somewhat picked up on saccade-related contributions in the signal, we don’t think such a contribution should be regarded as merely artefactual. Rather, it highlights the multimodal functional relevance of auditory alpha oscillatory activity and the naturally occurring interaction between audition and vision. In line with that, using a forward encoding procedure, Popov et al., (2021) nicely illustrate that in a purely auditory task, spatial tuning based on the hEOG signal is positively related to alpha tuning responses. Specifically, they argue that “auditory attention is linked to the visual system, at least in part, through pro-active orientation towards the relevant sound origin via saccades in the direction consistent with the sound origin” (Popov et al 2021, p.18).

### S5. Decoding analysis based on minimally preprocessed data

Artifact correction has been proposed to be less critical in decoding analysis, given that classifiers can – in principle – learn to ignore bad channels or suppress noise during training (Grootswagers, Wardle, & Carlson, 2017). Moreover, minimal preprocessing prevents unwanted artefacts and spurious decoding due to high-pass (van Driel, Olivers, & Fahrenfort, 2021) or low-pass (Grootswagers et al., 2017) filtering. Hence, the original decoding analysis was performed using only minimally preprocessed data. Following reviewer concerns, that artefacts might influence decoding performance, we modified the preprocessing pipeline for decoding to include artefact correction. The latter is now presented in the main manuscript. For reasons of transparency, here, we report the decoding results based on minimally preprocessed data. The latter yielded very comparable results.

The continuous data was epoched to create single-trial segments, ranging from −1000 to 4500 ms relative to cue onset, and baseline corrected (i.e., using the 200 ms time-period prior to cue onset as a baseline). Further, target-absent trials, incorrectly answered trials as well as trials with a response time < 200 (i.e., premature responses) were excluded. No filtering, trial rejection or ICA-based artefact correction was applied. The decoding analysis was performed as described in the main manuscript, except that five rather than three cross-validations were performed (accounting for the higher number of trials because no further trial rejection procedure was applied).

**Figure 5.**
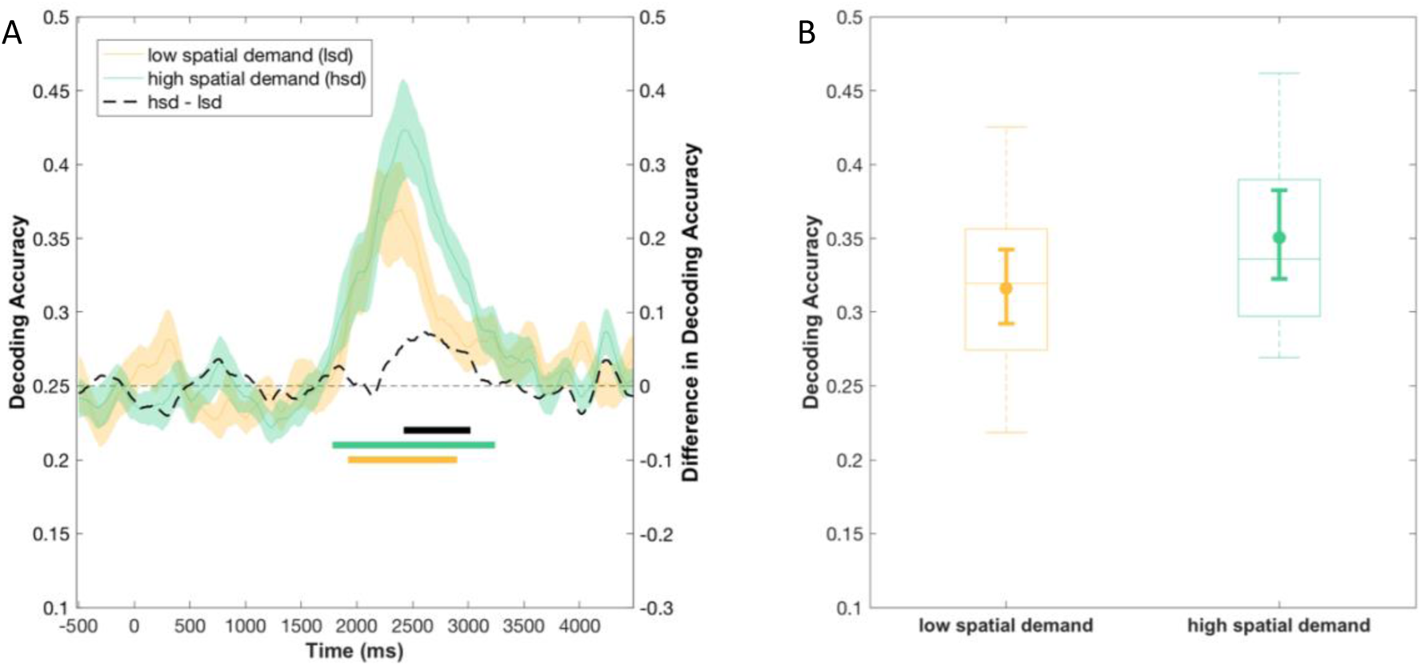
Location decoding based on the multivariate scalp distribution of alpha power (minimally preprocessed data. (A) Time-course of the average decoding accuracy results in the low (yellow) and high (green) spatial demand condition, respectively. The colored shading indicates ±1 SEM. Chance-level performance (i.e., 25%) is indicated by the grey dashed horizontal line. The yellow and green solid bars indicate significant decoding of the target location in the low and high spatial demand condition, respectively. The black solid bar denotes significant differences in decoding ability between the low and the high spatial demand condition. Note that only time-points in-between 1600 – 3800 ms were considered in the statistical analysis. (B) Boxplots refer to the average decoding accuracy in-between 1800 – 3200 ms relative to cue-onset (i.e., 200 – 1600 ms following sound array onset). As per convention, boxplots illustrate the interquartile range and the median. Whiskers extent to the 1.5 times the interquartile range. The superimposed circles show the average decoding accuracy, while the corresponding error bars denote the 95% bootstrap confidence interval of the mean (number of bootstrap samples = 10000).

Figure 5A shows the time-course of decoding accuracy for the low vs. high spatial demand condition, when decoding the exact target sound location based on the topography of alpha-band activity, as well as the difference in decoding accuracy between conditions. Decoding accuracy starts to rise above chance level (i.e., 25%) at around 1800 ms (i.e., 200 ms following sound array onset) and at first, increases continuously in both spatial demand conditions. In the high spatial demand condition, decoding accuracy reaches a peak at around 2180 ms (i.e., 580 ms post-sound onset), remains at this level for a couple hundred milliseconds and then gradually decreases throughout the remainder of the response interval; in the low spatial demand condition, decoding accuracy continues to rise beyond the peak in the high spatial demand condition until around ~2440 ms (i.e., 840 ms post-sound onset), and declines quite immediately thereafter, although it remains on a higher level compared to the low spatial demand condition. Toward the end of the response interval (i.e., around 3800 ms), decoding accuracy returns to chance level in both conditions. The cluster mass test revealed that decoding was significantly greater than chance in both spatial demand conditions. We identified a significant cluster following sound array onset in each of the two conditions (*p* < 10^−4^, see Figure 5A, solid green and yellow lines). In the high spatial demand condition, the cluster extends from around 1800 ms to ~3200 ms; in the low spatial demand condition, the cluster spans a time period in-between ~1900 ms and 2880 ms relative to sound array onset. Note, however, that cluster-based permutation test results should not be used to derive conclusions about the specific onset or offset of a certain effect (Sassenhagen & Draschkow, 2019).

The black, dashed line in Figure 5A illustrates the difference in decoding accuracy between the two spatial demand conditions. A cluster-corrected sign-permutation test indicated significant differences in decoding ability (*p* < .01, one-sided test, cluster extending from ~2440 – 3000 ms), with higher decoding accuracy in the high spatial demand condition compared to the low spatial demand condition.

Finally, we assessed the overall difference in decoding ability within the post-stimulus period (specifically, within the approximate time-window that resulted in above-chance decoding accuracy within both spatial demand conditions). A one-sided permutation test of the average decoding accuracy between 1800 – 3200 ms consistently revealed a significant difference in decoding accuracy between the spatial demand conditions (*p* = .001).

## Notes

1 Please note that the precise time window spans 415 ms. That is because the boundaries of the analysis time window refer to the actual sampling points that were present in the data: We first identified the peak in the grand average contralateral minus ipsilateral difference waveform. Then, the lower boundary of the analysis time window was determined by subtracting 200 ms from the peak latency and finding the sampling point that is closest to this value. Analogously, the upper boundary of the analysis time window was determined by adding 200 ms to the peak latency and finding the sampling point that is closest to this value.

2 Please note that the approach to compute alpha power for the decoding analysis (i.e., Hilbert transform) differs from the methods used for the univariate analyses of alpha power (i.e., Morlet wavelet convolution). For the univariate analyses, we wanted to keep the approach consistent with and comparable to our previous manuscripts (Begau, Klatt, Wascher, Schneider, & Getzmann, 2021; Klatt et al., 2019, 2018a); for the decoding analysis, we adopted the analysis approach from Bae and Luck (2018) and thus, applied a Filter Hilbert transformation. However, conducting the univariate analyses of alpha power using a Hilbert transform approach, essentially reproduces all results.

3 We thank two anonymous reviewers for suggesting the response-locked analyses.

4 Please note that the precise time window spans 406 ms. That is because the boundaries of the analysis time window refer to the actual sampling points that were present in the data: We first identified the peak in the grand average contralateral minus ipsilateral difference waveform. Then, the lower boundary of the analysis time window was determined by subtracting 200 ms from the peak latency and finding the sampling point that is closest to this value. Analogously, the upper boundary of the analysis time window was determined by adding 200 ms to the peak latency and finding the sampling point that is closest to this value

## Notes

### Competing Interest Statement

The authors have declared no competing interest.

### Summary of Updates

Major revisions concern: additional analysis of response-locked alpha-band ERSPs are included, further limitations are addressed in the discussion

## References

1. Adam, K. C. S., Vogel, E. K., & Awh, E. (2020). Multivariate analysis reveals a generalizable human electrophysiological signature of working memory load. Psychophysiology, 57(12), 1–17. https://doi.org/10.1111/psyp.13691

2. Anton-Erxleben, K., & Carrasco, M. (2013, March). Attentional enhancement of spatial resolution: Linking behavioural and neurophysiological evidence. Nature Reviews Neuroscience, Vol. 14, pp. 188–200. https://doi.org/10.1038/nrn3443

3. Arnau, S., Löffler, C., Rummel, J., Hagemann, D., Wascher, E., & Schubert, A. L. (2020). Inter-trial alpha power indicates mind wandering. Psychophysiology, 57(6), 1–14. https://doi.org/10.1111/psyp.13581

4. Bacigalupo, F., & Luck, S. J. (2019). Lateralized suppression of alpha-band EEG activity as a mechanism of target processing. Journal of Neuroscience, 39(5), 900–917. https://doi.org/10.1523/JNEUROSCI.0183-18.2018

5. Bae, G.-Y., & Luck, S. J. (2018). Dissociable Decoding of Spatial Attention and Working Memory from EEG Oscillations and Sustained Potentials. The Journal of Neuroscience, 38(2), 409–422. https://doi.org/10.1523/JNEUROSCI.2860-17.2017

6. Bae, G.-Y., & Luck, S. J. (2019). Appropriate Correction for Multiple Comparisons in Decoding of ERP Data: A Re-Analysis of Bae & Luck (2018). BioRxiv. https://doi.org/10.1101/672741

7. Bahramisharif, A., Heskes, T., Jensen, O., & van Gerven, M. A. J. (2011). Lateralized responses during covert attention are modulated by target eccentricity. Neuroscience Letters, 491, 35–39. https://doi.org/10.1016/j.neulet.2011.01.003

8. Bakdash, J. Z., & Marusich, L. R. (2017). Repeated measures correlation. Frontiers in Psychology, 8(MAR), 1–13. https://doi.org/10.3389/fpsyg.2017.00456

9. Banerjee, S., Snyder, A. C., Molholm, S., & Foxe, J. J. (2011). Oscillatory alpha-band mechanisms and the deployment of spatial attention to anticipated auditory and visual target locations: Supramodal or sensory-specific control mechanisms? The Journal of Neuroscience, 31(27), 9923–9932. https://doi.org/10.1523/JNEUROSCI.4660-10.2011

10. Begau, A., Klatt, L. I., Wascher, E., Schneider, D., & Getzmann, S. (2021). Congruent lip movements facilitate speech processing in a dynamic audiovisual multi-talker scenario: An ERP study with older and younger adults. Behavioural Brain Research, 412, 113436. https://doi.org/10.1101/2020.11.06.370841

11. Bigdely-Shamlo, N., Mullen, T., Kothe, C., Su, K.-M., & Robbins, K. A. (2015). The PREP pipeline: standardized preprocessing for large-scale EEG analysis. Frontiers in Neuroinformatics, 9(June), 1–20. https://doi.org/10.3389/fninf.2015.00016

12. Brungart, D. S., & Simpson, B. (2007). Cocktail party listening in a dynamic multitalker environment. Perception & Psychophysics, 69(1), 79–91.

13. Brungart, D. S., Simpson, B. D., Ericson, M. A., & Scott, K. R. (2001). Informational and energetic masking effects in the perception of multiple simultaneous talkers. The Journal of the Acoustical Society of America, 110(5), 2527–2538. https://doi.org/10.1121/1.1408946

14. Campbell, J. I. D., & Thompson, V. A. (2012). MorePower 6.0 for ANOVA with relational confidence intervals and Bayesian analysis. Behavior Research Methods, 44(4), 1255– 1265. https://doi.org/10.3758/s13428-012-0186-0

15. Carrasco, M., & McElree, B. (2001). Covert attention accelerates the rate of visual information processing. Proceedings of the National Academy of Sciences of the United States of America, 98(9), 5363–5367. https://doi.org/10.1073/pnas.081074098

16. Carrasco, M., Penpeci-Talgar, C., & Eckstein, M. (2000). Spatial covert attention increases contrast sensitivity across the CSF: Support for signal enhancement. Vision Research, 40(10–12), 1203–1215. https://doi.org/10.1016/S0042-6989(00)00024-9

17. Delorme, A., & Makeig, S. (2004). EEGLAB: an open source toolbox for analysis of single-trial EEG dynamics including independent component analysis. Journal of Neuroscience Methods, 134(2004), 9–21. https://doi.org/10.1016/j.jneumeth.2003.10.009

18. Deng, Y., Choi, I., & Shinn-Cunningham, B. G. (2020). Topographic specificity of alpha power during auditory spatial attention. NeuroImage, 207, 116360. https://doi.org/10.1016/j.neuroimage.2019.116360

19. Dietterich, T. G., & Balkiri, G. (1995). Solving Multiclass Learning Problems via Error-Correcting Output Codes. Journal of Artifical Intelligence Research, 2, 263–286.

20. Ericson, M. A., Brungart, D. S., & Simpson, B. D. (2004). Factors That Influence Intelligibility in Multitalker Speech Displays. In The International Journal of Aviation Psychology (Vol. 14). https://doi.org/10.1207/s15327108ijap1403

21. Feldmann-Wüstefeld, T., & Awh, E. (2019). Alpha-band activity tracks the zoom lens of attention. Journal of Cognitive Neuroscience, 32(2), 272–282. https://doi.org/10.1162/jocn_a_01484

22. Foster, J. J., Sutterer, D. W., Serences, J. T., Vogel, E. K., & Awh, E. (2017). Alpha-band oscillations enable spatially and temporally resolved tracking of covert spatial attention. Psychological Science, 28(7), 929–941. https://doi.org/10.1177/0956797617699167

23. Foxe, J. J., Simpson, G. V., & Ahlfors, S. P. (1998). Parieto-occipital ~10 Hz activity reflects anticipatory state of visual attention mechanisms. NeuroReport, 9(17), 3929–3933. https://doi.org/10.1097/00001756-199812010-00030

24. Fukuda, K., Mance, I., & Vogel, E. K. (2015). Α Power Modulation and Event-Related Slow Wave Provide Dissociable Correlates of Visual Working Memory. Journal of Neuroscience, 35(41), 14009–14016. https://doi.org/10.1523/JNEUROSCI.5003-14.2015

25. Gamble, M. L., & Luck, S. J. (2011). N2ac: An ERP component associated with the focusing of attention within an auditory scene. Psychophysiology, 48(8), 1057–1068. https://doi.org/10.1111/j.1469-8986.2010.01172.x

26. Getzmann, S., Klatt, L. I., Schneider, D., Begau, A., & Wascher, E. (2020). EEG correlates of spatial shifts of attention in a dynamic multi-talker speech perception scenario in younger and older adults. Hearing Research, 398, 108077. https://doi.org/10.1016/j.heares.2020.108077

27. Hanslmayr, S., Spitzer, B., & Bäuml, K. H. (2009). Brain oscillations dissociate between semantic and nonsemantic encoding of episodic memories. Cerebral Cortex, 19(7), 1631–1640. https://doi.org/10.1093/cercor/bhn197

28. Hebart, M. N., & Baker, C. I. (2017). Deconstructing multivariate decoding for the study of brain function. NeuroImage, (2017). https://doi.org/10.1016/j.neuroimage.2017.08.005

29. Hentschke, H., & Stüttgen, M. C. (2011). Computation of measures of effect size for neuroscience data sets. European Journal of Neuroscience, 34, 1887–1894. https://doi.org/10.1111/j.1460-9568.2011.07902.x

30. Ikkai, A., Dandekar, S., & Curtis, C. E. (2016). Lateralization in alpha-band oscillations predicts the locus and spatial distribution of attention. PloS One, 11(5), e0154796. https://doi.org/10.1371/journal.pone.0154796

31. Kiesel, A., Miller, J., Jolicœur, P., & Brisson, B. (2008). Measurement of ERP latency differences: A comparison of single-participant and jackknife-based scoring methods. Psychophysiology, 45, 250–274. https://doi.org/10.1111/j.1469-8986.2007.00618.x

32. Klatt, L.-I., Getzmann, S., Begau, A., & Schneider, D. (2019). A dual mechanism underlying retroactive shifts of auditory spatial attention: dissociating target- and distractor-related modulations of alpha lateralization. Scientific Reports, 10, 13860. https://doi.org/10.1101/2019.12.19.882324

33. Klatt, L.-I., Getzmann, S., Wascher, E., & Schneider, D. (2018a). Searching for auditory targets in external space and in working memory: Electrophysiological mechanisms underlying perceptual and retroactive spatial attention. Behavioural Brain Research, 353, 98–107. https://doi.org/10.1016/j.bbr.2018.06.022

34. Klatt, L.-I., Getzmann, S., Wascher, E., & Schneider, D. (2018b). The contribution of selective spatial attention to sound detection and sound localization: Evidence from event-related potentials and lateralized alpha oscillations. Biological Psychology, 138, 133–145. https://doi.org/10.1016/j.biopsycho.2018.08.019

35. Krause, C. M., Sillanmaki, L., Koivisto, M., Saarela, C., Haggqvist, A., Laine, M., & Hamalainen, H. (2000). The effects of memory load on event-related (EEG) desynchronisation and synchronisation. Clinical Neurophysiology, 111, 2071–2078. https://doi.org/10.1016/S1388-2457(00)00429-6

36. Lee, C.-C., & Middlebrooks, J. C. (2011). Auditory Cortex Spatial Sensitivity Sharpens During Task Performance. Nature Neuroscience, 14(1), 108–114. https://doi.org/doi:10.1038/nn.2713.

37. Liesefeld, H. R. (2018). Estimating the Timing of Cognitive Operations With MEG / EEG Latency Measures : A Primer, a Brief Tutorial, and an Implementation of Various Methods. Frontiers in Neuroscience, 12(October), 1–11. https://doi.org/10.3389/fnins.2018.00765

38. Luck, S. J. (2014). An introduction to the event-related potential technique (2nd ed.). MIT Press.

39. Luck, S. J., & Gaspelin, N. (2017). How to get statistically significant effects in any ERP experiment (and why you shouldn’t). Psychophysiology, 54(1), 146–157. https://doi.org/10.1111/psyp.12639

40. Miller, J., Patterson, T., & Ulrich, R. (1998). Jackknife-based method for measuring LRP onset latency differences. Psychophysiology, 35(1), 99–115. https://doi.org/10.1111/1469-8986.3510099

41. Murphy, S., Spence, C. J., & Dalton, P. (2017). Auditory perceptual load: A review. Hearing Research, 352, 40–48. https://doi.org/10.1016/j.heares.2017.02.005

42. Oldfield, R. C. (1971). The assessment and analysis of handedness: the Edinburgh inventory. Neuropsychologia, 9, 97–113. Retrieved from http://www.ncbi.nlm.nih.gov/pubmed/5146491

43. Perrin, F., Pernier, J., Bertrand, O., & Echallier, J. F. (1989). Spherical splines for scalp potential and current density mapping. Electroencephalography and Clinical Neurophysiology, 72(2), 184–187. https://doi.org/10.1016/0013-4694(89)90180-6

44. Pion-Tonachini, L., Kreutz-Delgado, K., & Makeig, S. (2019). ICLabel: An automated electroenceophalographic independent component classifier, dataset, and website. NeuroImage, 198, 181–197. https://doi.org/10.1016/j.neuroimage.2019.05.026

45. Poch, C., Capilla, A., Hinojosa, J. A., & Campo, P. (2017). Selection within working memory based on a color retro-cue modulates alpha oscillations. Neuropsychologia, 106, 133–137. https://doi.org/10.1016/j.neuropsychologia.2017.09.027

46. Popov, T., Gips, B., Kastner, S., & Jensen, O. (2019). Spatial specificity of alpha oscillations in the human visual system. Human Brain Mapping, (June), 1–9. https://doi.org/10.1002/hbm.24712

47. Popov, T., Gips, B., Weisz, N., & Jensen, O. (2021). Brain areas associated with visual spatial attention display topographic organization during auditory spatial attention. BioRxiv. https://doi.org/https://doi.org/10.1101/2021.03.15.435371

48. Reiser, J. E., Wascher, E., Rinkenauer, G., & Arnau, S. (2020). Cognitive-motor interference in the wild: Assessing the effects of movement complexity on task switching using mobile EEG. European Journal of Neuroscience. https://doi.org/10.1111/ejn.14959

49. Richardson, J. T. E. (2011). Eta squared and partial eta squared as measures of effect size in educational research (2nd ed.). Educational Research Review, 6(2), 135–147. https://doi.org/10.1016/j.edurev.2010.12.001

50. Rihs, T. A., Michel, C. M., & Thut, G. (2007). Mechanisms of selective inhibition in visual spatial attention are indexed by α-band EEG synchronization. European Journal of Neuroscience, 25(2), 603–610. https://doi.org/10.1111/j.1460-9568.2007.05278.x

51. Rousselet, G. A. (2012). Does filtering preclude us from studying ERP time-courses? Frontiers in Psychology, 3(MAY), 1–9. https://doi.org/10.3389/fpsyg.2012.00131

52. Sassenhagen, J., & Draschkow, D. (2019). Cluster-based permutation tests of MEG/EEG data do not establish significance of effect latency or location. Psychophysiology, 56(6), e13335. https://doi.org/10.1111/psyp.13335

53. Schneider, D., Göddertz, A., Haase, H., Hickey, C., & Wascher, E. (2019). Hemispheric asymmetries in EEG alpha oscillations indicate active inhibition during attentional orienting within working. Behavioural Brain Research, 359, 38–46. https://doi.org/10.1016/j.bbr.2018.10.020

54. Schneider, D., Mertes, C., & Wascher, E. (2016). The time course of visuo-spatial working memory updating revealed by a retro-cuing paradigm. Scientific Reports, 6, 21442. https://doi.org/10.1038/srep21442

55. Shiell, M. M., Hausfeld, L., & Formisano, E. (2018). Activity in human auditory cortex represents spatial separation between concurrent sounds. The Journal of Neuroscience, 38(21), 4977–4984. https://doi.org/10.1523/JNEUROSCI.3323-17.2018

56. Soper, D. S. (2020). p-Value Calculator for an F-Test [Software]. Retrieved December 16, 2020, from https://www.danielsoper.com/statcalc

57. Sprague, T. C., Saproo, S., & Serences, J. T. (2015). Visual attention mitigates information loss in small- and large scale neural codes. Trends in Cognitive Sciences, 19(4), 215–226. https://doi.org/10.1016/j.tics.2015.02.005

58. van der Heijden, K., Rauschecker, J. P., Formisano, E., Valente, G., & de Gelder, B. (2018). Active sound localization sharpens spatial tuning in human primary auditory cortex. Journal of Neuroscience, 38(40), 8574–8587. https://doi.org/10.1523/JNEUROSCI.0587-18.2018

59. VanRullen, R. (2011). Four common conceptual fallacies in mapping the time course of recognition. Frontiers in Psychology, 2(DEC), 1–6. https://doi.org/10.3389/fpsyg.2011.00365

60. Voytek, B., Samaha, J., Rolle, C. E., Greenberg, Z., Gill, N., Porat, S., … Gazzaley, A. (2017). Preparatory encoding of the fine scale of human spatial attention. Journal of Cognitive Neuroscience, 29(7), 1302–1310. https://doi.org/10.1162/jocn_a_01124

61. Wang, S., Megla, E. E., & Woodman, G. F. (2021). Stimulus-induced alpha suppression tracks the difficulty of attentional selection, not visual working memory storage. Journal of Cognitive Neuroscience, 33(3), 536–562. https://doi.org/10.1162/jocn_a_01637

62. Winkler, I., Debener, S., Muller, K. R., & Tangermann, M. (2015). On the influence of high-pass filtering on ICA-based artifact reduction in EEG-ERP. Proceedings of the Annual International Conference of the IEEE Engineering in Medicine and Biology Society, EMBS, 2015-Novem, 4101–4105. https://doi.org/10.1109/EMBC.2015.7319296

63. Winter, B. (2011). The F distribution and the basic principle behind ANOVAs. Retrieved from http://menzerath.phonetik.uni-frankfurt.de/teaching/R/bw_anova_general_HR.pdf

64. Wolff, M. J., Jochim, J., Akyürek, E. G., & Stokes, M. G. (2017). Dynamic hidden states underlying working-memory-guided behavior. Nature Neuroscience, 20(6), 864–871. https://doi.org/10.1038/nn.4546

65. Worden, M. S., Foxe, J. J., Wang, N., & Simpson, G. V. (2000). Anticipatory biasing of visuospatial attention indexed by retinotopically specific α-band electroencephalography increases over occipital cortex. The Journal of Neuroscience, 20(6), RC63. https://doi.org/10.1523/JNEUROSCI.20-06-j0002.2000

## References

66. Deng, Y., Choi, I., & Shinn-Cunningham, B. G. (2020). Topographic specificity of alpha power during auditory spatial attention. NeuroImage, 207, 116360. https://doi.org/10.1016/j.neuroimage.2019.116360

67. Grootswagers, T., Wardle, S. G., & Carlson, T. A. (2017). Decoding dynamic brain patterns from evoked responses: A tutorial on multivariate pattern analysis applied to time series neuroimaging data. Journal of Cognitive Neuroscience, 29(4), 677–697. https://doi.org/10.1162/jocn_a_01068

68. Klatt, L.-I., Getzmann, S., Wascher, E., & Schneider, D. (2018). Searching for auditory targets in external space and in working memory: Electrophysiological mechanisms underlying perceptual and retroactive spatial attention. Behavioural Brain Research, 353, 98–107. https://doi.org/10.1016/j.bbr.2018.06.022

69. Miller, J., Patterson, T., & Ulrich, R. (1998). Jackknife-based method for measuring LRP onset latency differences. Psychophysiology, 35(1), 99–115. https://doi.org/10.1111/1469-8986.3510099

70. Popov, T., Gips, B., Weisz, N., & Jensen, O. (2021). Brain areas associated with visual spatial attention display topographic organization during auditory spatial attention. BioRxiv. https://doi.org/https://doi.org/10.1101/2021.03.15.435371

71. Sassenhagen, J., & Draschkow, D. (2019). Cluster-based permutation tests of MEG/EEG data do not establish significance of effect latency or location. Psychophysiology, 56(6), e13335. https://doi.org/10.1111/psyp.13335

72. van Driel, J., Olivers, C. N. ., & Fahrenfort, J. J. (2021). High-pass filtering artifacts in multivariate classification of neural time series data. Journal of Neuroscience Methods, 352, 109080. https://doi.org/10.1101/530220

73. van Moorselaar, D., Foster, J. J., Sutterer, D. W., Theeuwes, J., Olivers, C. N. L., & Awh, E. (2018). Spatially Selective Alpha Oscillations Reveal Moment-by-Moment Trade-offs between Working Memory and Attention. Journal of Cognitive Neuroscience, 30(2), 256–266. https://doi.org/10.1162/jocn_a_01198

